# Agentic Discovery Transfer Enables Cross-Cancer Discovery at Scale

**DOI:** 10.64898/2026.10.28.762506

**Authors:** Lingxin Cheng, Jack Shireman, Akash Gogate, Mary M. Stangis, Yujia Cai, Elliot Xie, Parth Khatri, Mahua Dey, Christina Kendziorski

## Abstract

Findings from cancer studies have the potential to inform research across multiple cancer types, yet they are rarely evaluated beyond their original setting. We present SMART, a multi-agent large language model framework that autonomously evaluates whether findings from published cancer studies transfer to new cancer contexts, a task we term discovery transfer. Given a source manuscript and a target single-cell RNA-sequencing dataset, SMART extracts the manuscript’s findings, designs and executes analyses to test them, and assesses the extent to which each finding transfers. In manual audits, 97% of SMART’s outputs were correct. We applied SMART to 121 studies and six target cancer types, yielding 726 reports. Findings transferred in 46% of cross-cancer reports, revealing shared prognostic signatures, conserved and context-specific mechanisms, and potential therapeutic opportunities. These results demonstrate how agentic discovery transfer can extend published cancer findings beyond their original contexts and identify new avenues for biological and clinical investigation.

---

Cancer research generates tens of thousands of studies each year^1,2^, each representing a substantial investment of time, funding, and expertise. Although most studies focus on a particular cancer type, their findings may have implications far beyond the cancer in which they were made. HER2-targeted therapy, for example, first established in breast cancer, also improves survival in HER2-positive gastric cancer^3^; mismatch-repair deficiency, first characterized in colorectal cancer, is now known to predict response to immune checkpoint blockade across tumor types^4^.

Despite the potential for advances, findings are rarely evaluated systematically across cancers, as doing so requires substantial time, domain expertise, and computational resources. In particular, to determine whether findings reported in a given manuscript are relevant in another context, an investigator must read and interpret the source paper, identify its key claims, decide which of those claims are biologically and technically testable in the target setting, and design and execute the necessary analyses. This process, defined here as discovery transfer, is slow, labor-intensive, and inherently selective, as the findings chosen for follow-up are inevitably shaped by the investigator’s interests, expertise, and analytic capacity.

As a result of the resources required for discovery transfer, relatively few potentially transferable findings are systematically assessed, and the vast majority of insights published in the research literature remain underutilized. Consider the lung cancer prognostic signature ORACLE^5,6^, first reported in 2019. Of the 161 studies citing the ORACLE papers, to our knowledge only one has tested it in another cancer, finding it prognostic in pleural mesothelioma more than four years after ORACLE’s publication^7^. As we show here, its association with poor survival also extends to liver and pancreatic cancer.

Discovery transfer is especially valuable, and especially laborious, for single-cell RNA-sequencing (scRNA-seq) studies of cancer, whose findings concern specific cell populations, transcriptional programs, cell–cell communication networks, and functional pathways^8–10^. Such findings, already tested in their original setting, are strong starting points for hypotheses in a new cancer dataset.

While general-purpose agents can carry out open-ended research requests, including scRNA-seq analysis^11–17^, and/or autonomously generate and test new hypotheses^18^, none is designed to test whether reported findings transfer to an independent dataset. As illustrated here, general-purpose agents given this task replaced the source study’s validated definitions with their own and left some of its claims untested (Supplementary Notes 1 and 2, Supplementary Fig. 1, Supplementary Table 1 and Supplementary Data 1–6).

Here we present SMART, a <u>s</u>ystem of <u>m</u>ulti-<u>a</u>gent large language models for discovery transfer in scRNA-seq studies (Fig. 1). Given a published study (the source study) and an scRNA-seq dataset from another cancer (the target dataset), SMART extracts the study’s clinical and mechanistic findings, decides which can be tested, writes and runs the analyses, reports which findings transfer, and suggests follow-up. We validate SMART using a positive-control experiment, across repeated runs, and with decoy summaries in which every finding is reversed. In manual audits of the positive control and other transfer cases, 97% of SMART’s outputs were correct on average.

**Figure 1.**
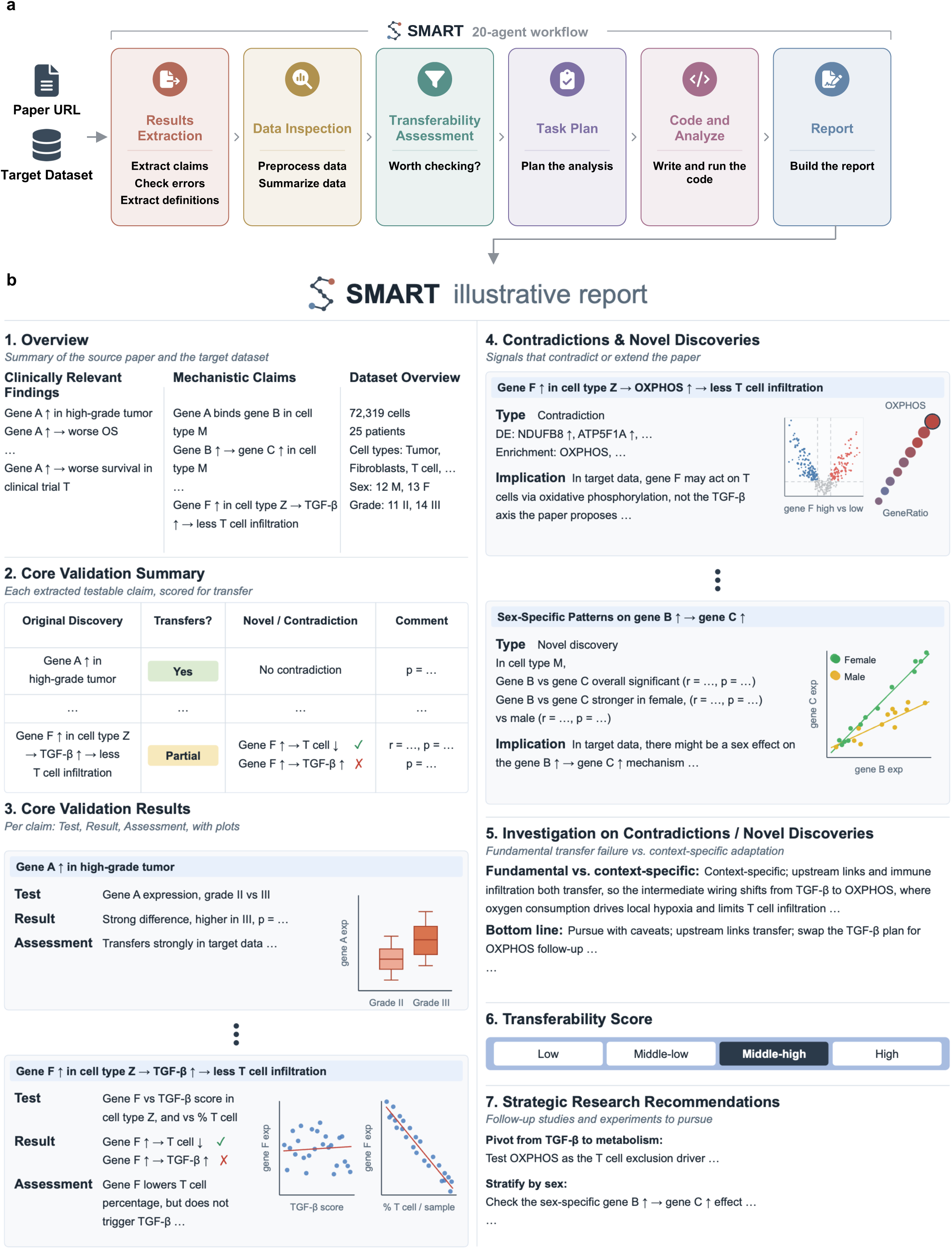
SMART is an agentic discovery transfer framework for evaluating whether results from published studies transfer to scRNA-seq datasets. A user provides two inputs: a source paper of interest (PubMed URL) and a target scRNA-seq dataset with its associated metadata. SMART processes these inputs through six sequential modules, consisting of 20 specialized agents in total, to identify the source paper’s discoveries and to evaluate which of those discoveries transfer to the target dataset. **(a)** The SMART workflow. The Results Extraction module reads the source paper and extracts its clinically relevant findings and mechanistic claims, then error-checks each against the original text and figures, and resolves the biological terms they depend on into operational definitions. The Data Inspection module prepares the target dataset and summarizes its cell types and metadata. The Transferability Assessment module checks whether the target dataset contains the cell types and transcriptional readouts that the source mechanism requires. The Task Plan module translates the extracted findings into an execution plan of analyses, with every parameter set from the paper and the dataset. The Code and Analyze module writes the plan as R scripts built from a curated library of single-cell analysis functions, then runs and debugs them. The Report module interprets every plot and statistic and compiles them into a structured report. Individual agents within each module are described in Methods. **(b)** Structure of a SMART report. To illustrate the report format independently of any particular biology, we deliberately use generic placeholders (genes A, B, C and F; cell types M and Z) in place of real gene and cell-type names; an example with real data is shown in Fig. 2, and full reports are provided in Supplementary Data 5 and 8–12. Section 1 of SMART’s report states the paper’s clinically relevant findings and mechanistic claims alongside a summary of the target dataset. Section 2 reports whether each testable claim transfers (Yes, Partial or No) in a single summary table. Section 3 expands each claim in detail, describing what was tested, the resulting statistics and an assessment of the result, paired with the supporting figures. Section 4 reports results that contradict the source paper and new signals in the target dataset that the source paper did not report. Section 5 investigates whether each contradiction reflects a fundamental failure of the claim’s logic or a context-specific adaptation. Section 6 assigns an overall transferability score (Low, Middle-low, Middle-high or High), and Section 7 gives prioritized, conditional recommendations for follow-up analyses and experiments.

Evaluation of SMART across 121 studies and six cancer types demonstrates its ability to systematically assess whether published clinical and mechanistic findings transfer to new cancer contexts. Beyond identifying findings that are conserved across cancers, SMART reveals context-specific differences in biological mechanisms, extends prognostic signatures to new cancer types, and identifies potential therapeutic opportunities. By connecting transferable claims from separate studies, SMART also generates new mechanistic hypotheses, several of which are supported by independent evidence. Finally, the resulting collection of tested findings provides a searchable resource through which researchers can explore relationships across cancer types and generate new hypotheses.

## Results

### SMART transfers findings through a 20-agent workflow

SMART consists of six modules, each composed of specialized agents: Results Extraction, Data Inspection, Transferability Assessment, Task Plan, Code and Analyze, and Report (Fig. 1a and Methods). To run SMART, a user specifies the published study via a PubMed URL and supplies the target scRNA-seq dataset with its metadata (for example, as Seurat^19^, Scanpy^20^ or Cell Ranger^21^ output).

The Results Extraction module identifies the source study’s main results, organized into clinically relevant findings and mechanistic claims (Supplementary Note 2 and Supplementary Data 6). The Data Inspection module summarizes the cell types and metadata of the target dataset (Supplementary Note 3). The Transferability Assessment module acts as a gate, determining whether discovery transfer is feasible in the target dataset at all, for example whether the cell types central to the study are present. The Task Plan module then determines which individual findings and claims are testable and specifies analysis tasks for them, which the Code and Analyze module translates into executable R scripts, runs and debugs. Finally, the Report module presents plots, statistics and a verdict for each claim (supported, partially supported or not supported in the target dataset), highlights results that contradict or extend the source paper, and gives an overall transferability score (Low, Middle-low, Middle-high or High) with suggested follow-up (Fig. 1b). We consider a study’s findings to transfer when the score is Middle-high or High (Methods).

### SMART accurately recovers and evaluates published findings

We first paired a study with its own scRNA-seq data as a positive control. Because the study’s findings were mostly derived from this dataset, SMART should extract them correctly and find that they hold. We applied SMART to the colorectal cancer study by Qi et al.^22^, whose dataset consists of 54,103 cells from paired tumor and adjacent normal tissue from five patients. Qi et al. found that FAP⁺ fibroblasts and SPP1⁺ macrophages are enriched in colorectal tumors and linked to poor prognosis, and proposed that these cells interact and build an extracellular matrix (ECM) barrier that excludes lymphocytes from the tumor core (Fig. 2a).

**Figure 2.**
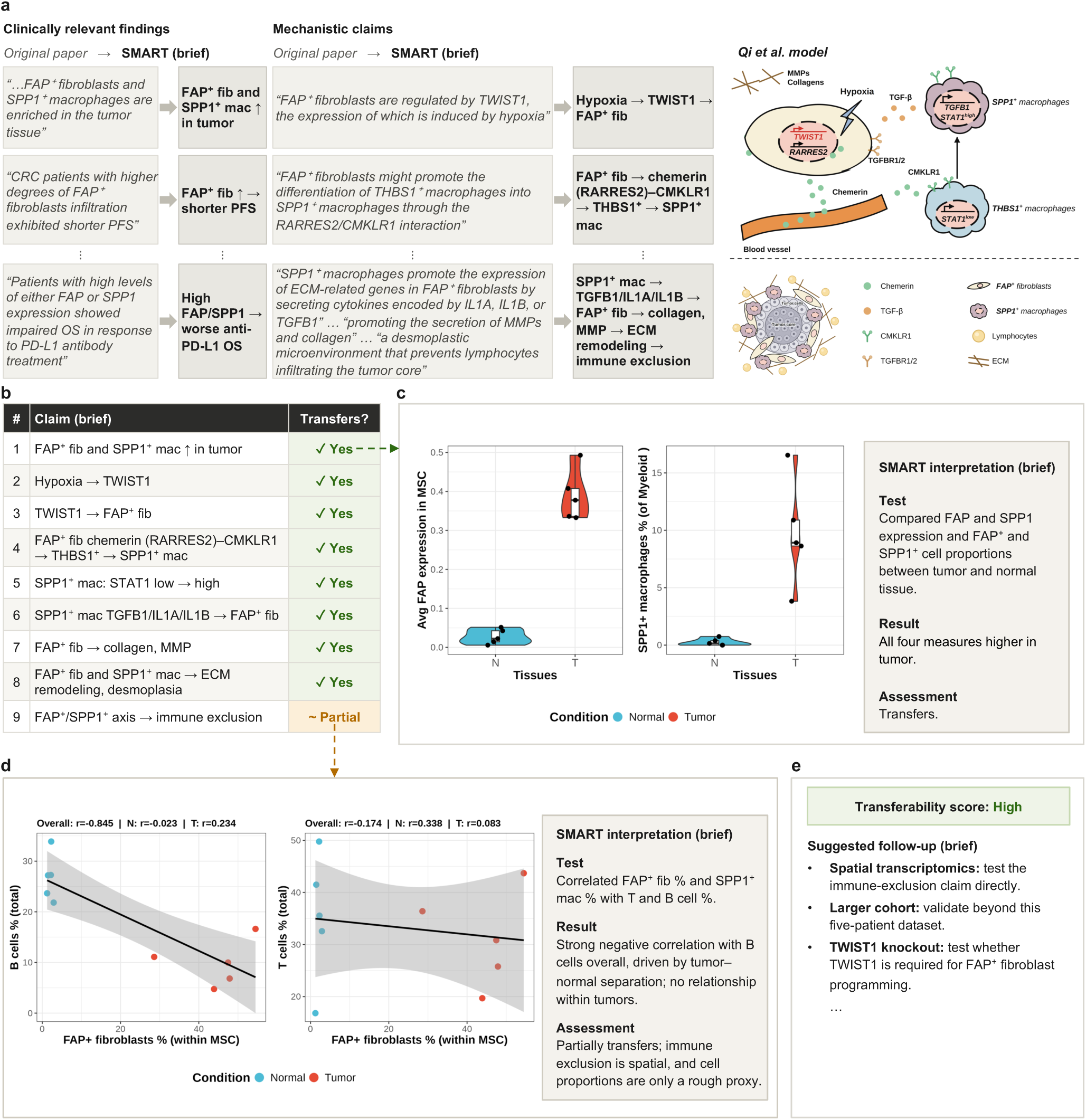
SMART reproduces a colorectal cancer study’s findings in the study’s own scRNA-seq data. We applied SMART to the colorectal cancer study by Qi et al.^22^, supplying the PubMed URL for the manuscript and the study’s own scRNA-seq dataset as a positive control. The full report is provided in Supplementary Data 8. **(a)** Selected clinically relevant findings (left) and mechanistic claims (middle) from the original text of the source paper are shown beside SMART’s extracted output (abbreviated for display); the full extracted text is given in Supplementary Data 7. The right panel shows the cross-talk model proposed by Qi et al. (redrawn and rearranged from Fig. 8j in ref. 22). SMART’s extracted claims map onto every node of the Qi et al. model. **(b)** SMART decomposed the extracted findings into nine testable claims, then wrote and executed the code to evaluate each. Eight claims were found to transfer to the target dataset and one, immune exclusion, to transfer partially. One example is shown in panel c. **(c)** Claim 1 in panel b. SMART’s plots of *FAP* expression in mesenchymal stromal cells (left) and SPP1⁺ macrophages as a percentage of myeloid cells (middle) in normal and tumor tissue from the five patients, and SMART’s written output (brief version, right). **(d)** The immune-exclusion claim (claim 9 in panel b), which transferred partially. SMART’s plots of B cell (left) and T cell (middle) proportions against FAP⁺ fibroblast proportions, with samples colored by tissue, and SMART’s written output (brief version, right). **(e)** SMART’s overall transferability score for the study, together with the follow-up experiments it proposed (abbreviated). CRC, colorectal cancer; PFS, progression-free survival; OS, overall survival; ECM, extracellular matrix; fib, fibroblasts; mac, macrophages; MMP, matrix metalloproteinase; N, normal; T, tumor.

SMART accurately recovered the major clinical and mechanistic findings, as verified against the source manuscript (Fig. 2a and Supplementary Data 7). It decomposed them into nine testable claims, and eight of them transferred (Fig. 2b,c). The remaining claim, that the ECM barrier excludes lymphocytes, is about spatial organization, which dissociated scRNA-seq data cannot fully capture. SMART flagged this limitation and used correlations of FAP⁺ fibroblast and SPP1⁺ macrophage proportions with T and B cell proportions as a proxy. These cells were negatively correlated with B cells across all samples, but this reflected the separation between tumor and normal tissue, and within tumors neither T nor B cell proportions were strongly related to them, so SMART judged the claim to transfer partially (Fig. 2d). Overall, SMART assigned the study a High transferability score and suggested follow-up experiments, such as spatial transcriptomics to test immune exclusion directly and validation in more patients, given that the dataset includes only five (Fig. 2e and Supplementary Data 8).

We manually validated SMART in 10 cases in total: the positive control above and 9 discovery transfer cases (Supplementary Note 4 and Supplementary Table 2). Averaged across all audited steps, from finding extraction to the testing and reporting of each claim, 97% of audited outputs were correct, and the outputs of the Data Inspection and Transferability Assessment modules were correct in all cases (Supplementary Data 7). Across three independent runs of each case, the transfer decision (transfer versus no transfer) was identical in all 10 cases, and the transferability score differed by at most one level (Supplementary Table 3). To test whether SMART simply confirms the claims it is given, we reran 15 of the reports with reversed source summaries as decoys (Methods). All 15 decoy reports were scored Middle-low or Low, including the 12 whose original reports were scored Middle-high or High (Supplementary Table 4).

### SMART evaluates 121 studies across six cancers

We next applied SMART at scale to 121 scRNA-seq studies of glioblastoma or colorectal, liver, lung, ovarian or pancreatic cancer, published in Nature Portfolio journals during 2024–2025, running each against six target datasets, one per cancer type, to generate 726 end-to-end reports (Fig. 3a and Supplementary Table 5). Of the 605 reports in which the source and target cancers differed, 46% transferred, ranging from 17% to 67% across pairs of cancers, compared with 75– 88% when they were the same.

**Figure 3.**
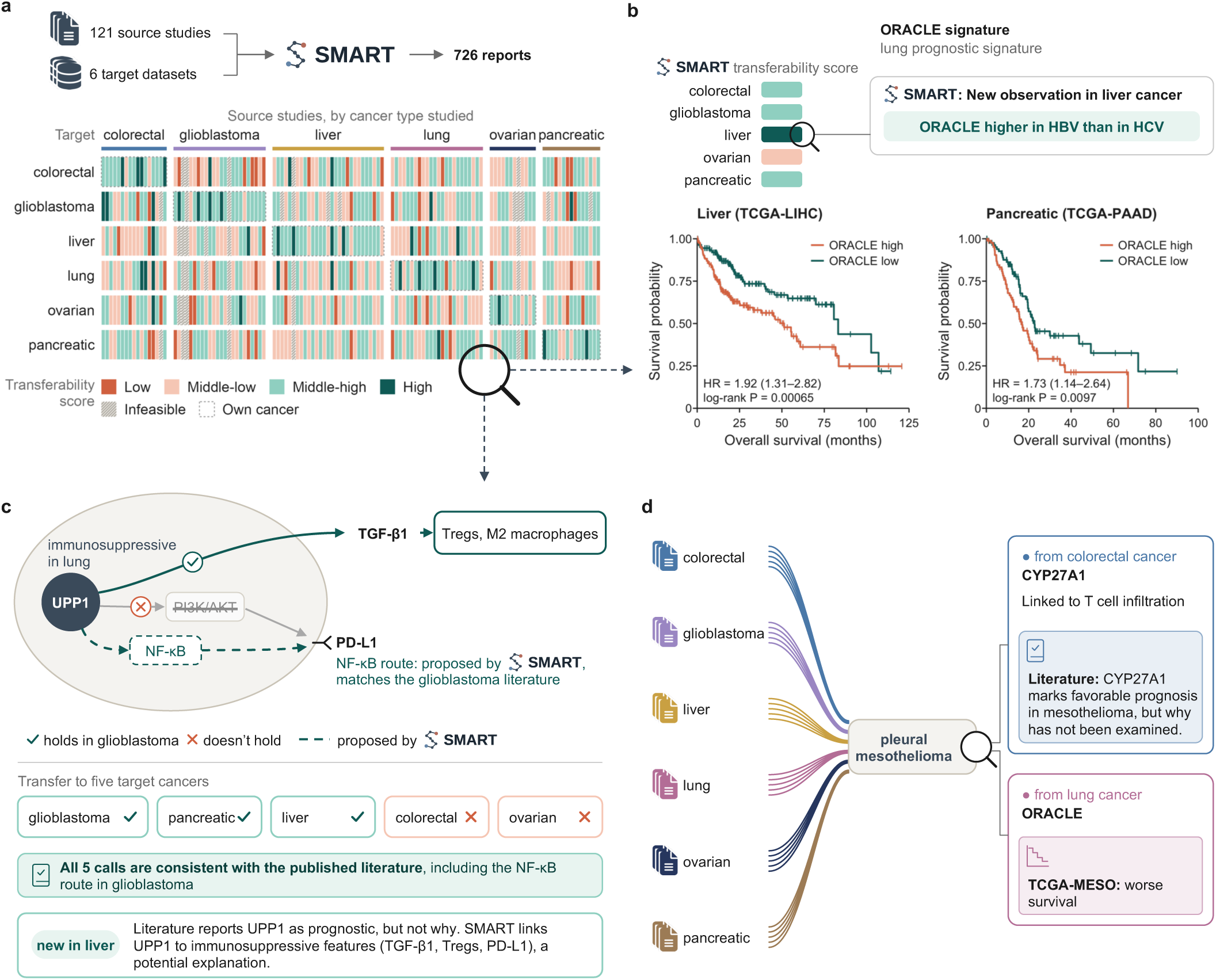
Applying SMART at scale identifies published discoveries that transfer beyond the cancer type in which they were made. We applied SMART to every pairing of the 121 source studies and the six target datasets. **(a)** The cross-cancer transfer landscape derived from the 726 reports. Each column represents one source study and each row one target dataset; columns are grouped by the cancer in which the source study was performed; dashed outlines mark pairs in which the target dataset is from the same cancer type as the source study; cells marked Infeasible are pairs that the Transferability Assessment module stopped before analysis, as described in Fig. 1a. Each cell is colored by SMART’s transferability score (Low, Middle-low, Middle-high or High). **(b)** ORACLE, a lung cancer prognostic signature, transfers to most other cancers tested. Top, SMART’s transferability score for ORACLE in each non-lung target, colored as in panel a, and an observation SMART reported in liver cancer: higher ORACLE scores in hepatitis B virus (HBV)- than in hepatitis C virus (HCV)-associated tumors. Bottom, overall survival in TCGA liver (TCGA-LIHC, n = 336) and pancreatic (TCGA-PAAD, n = 171) cohorts for patients with ORACLE scores above or below the median; hazard ratios (95% confidence intervals) and log-rank P values are given in each panel. Associations in the other three cohorts were not significant (Supplementary Note 5 and Supplementary Fig. 2). **(c)** Transfer of findings from a lung cancer study of *UPP1*-driven immunosuppression^23^ to five target cancers. Top, the lung mechanism tested in glioblastoma (check mark, holds; cross, does not hold), with the NF-κB route proposed by SMART in place of PI3K/AKT (dashed). Middle, SMART’s transfer call for each target cancer, all five of which are consistent with the published literature. Bottom, a potential explanation suggested by SMART for the reported prognostic role of *UPP1* in liver cancer^27^. Full reports are provided in Supplementary Data 10. **(d)** Transfer of findings from well-studied cancers to a rare one. SMART evaluated transfer from all 121 source studies to a pleural mesothelioma scRNA-seq dataset (lines are schematic and colored by source cancer). Right, two transferring findings. Top, *CYP27A1*, from a colorectal cancer study^30^: SMART linked it to T cell infiltration in epithelioid tumors, and the literature reports it as a favorable prognostic marker in mesothelioma^31^. Bottom, the lung prognostic signature ORACLE, which we found to be associated with worse survival in TCGA-MESO (n = 69; hazard ratio = 3.04, 95% confidence interval 1.78–5.18; log-rank P = 2.2 × 10⁻⁵). Full reports are in Supplementary Data 11 and the survival analysis in Supplementary Note 5.

### SMART finds a lung signature prognostic in other cancers

Of the six source studies reporting clinically oriented signatures or classifiers (prognostic, predictive or diagnostic), one transferred most widely, to four of the five other cancers. That study^5^ prospectively validated ORACLE^6^, a 23-gene prognostic signature whose high scores are associated with poor survival in lung cancer. The signature was also shown to be associated with chromosomal instability and to increase with disease progression. In liver cancer, for example, SMART found that ORACLE-high hepatocytes were enriched for mitotic processes such as chromosome segregation, and ORACLE scores increased from normal tissue to tumor to portal vein tumor thrombus, mirroring the pattern reported in lung cancer. SMART also observed higher ORACLE scores in hepatitis B virus- than in hepatitis C virus-associated tumors. ORACLE also transferred to colorectal and pancreatic cancer and glioblastoma, but not to ovarian cancer (Fig. 3b and Supplementary Data 9).

In most reports, SMART recommended further testing ORACLE against survival data (Supplementary Data 9). We therefore followed up in bulk RNA-sequencing cohorts from The Cancer Genome Atlas (TCGA) and found that higher ORACLE scores were significantly associated with poorer survival in liver cancer (hazard ratio [HR] = 1.92, 95% confidence interval [CI] 1.31–2.82, P = 0.00065) and pancreatic cancer (HR = 1.73, 95% CI 1.14–2.64, P = 0.0097; Fig. 3b), but not in ovarian cancer, glioblastoma or colorectal cancer (Supplementary Note 5 and Supplementary Fig. 2). Because proliferation is associated with survival in many cancers, we also compared ORACLE with random 23-gene signatures matched to ORACLE’s correlation with proliferation. ORACLE was more strongly associated with survival than 98% of these signatures in liver cancer (empirical P = 0.021) and 94% in pancreatic cancer (P = 0.062) (Supplementary Note 5).

### SMART extends an immunosuppressive mechanism across cancers

Among findings on the tumor microenvironment, a lung cancer study^23^ reported that *UPP1*-high tumor cells drive immunosuppression by releasing cytokines, most prominently TGF-β1, which shift surrounding cells toward immunosuppressive states such as regulatory T cells and M2-like macrophages. *UPP1* also activates PI3K/AKT/mTOR signaling, which upregulates PD-L1 on tumor cells. SMART reported that the *UPP1* findings transferred to glioblastoma, pancreatic cancer and liver cancer, but not to colorectal or ovarian cancer (Fig. 3c and Supplementary Data 10).

In glioblastoma, *UPP1*-high glioma cells expressed more *TGFB1*, and *UPP1* expression correlated with regulatory T cell infiltration and M2-like macrophage polarization. *UPP1* was not correlated with PI3K/AKT pathway scores but remained strongly correlated with PD-L1 expression in primary glioblastoma. Because genes upregulated in *UPP1*-high glioma cells were enriched for NF-κB signaling, SMART proposed NF-κB as a candidate route from *UPP1* to PD-L1 in place of PI3K/AKT. This is consistent with glioma and glioblastoma studies linking *UPP1* to poor survival, regulatory T cell infiltration and immune checkpoint expression^24,25^ and proposing that it activates NF-κB signaling^25^. SMART similarly found *UPP1* associated with immunosuppressive cytokines and M2-like macrophages in pancreatic cancer, where *UPP1* has been shown to support tumor growth and is associated with poor survival^26^. The absence of transfer is also consistent with prior reports that *UPP1* is downregulated in colorectal tumors^25^ and has no prognostic association in ovarian cancer^27^.

In liver cancer, SMART found that *UPP1* correlated with immunosuppressive cytokine expression in hepatocytes, that *UPP1*-high samples showed greater regulatory T cell infiltration, and that *UPP1* was associated with higher PI3K/AKT/mTOR pathway activity and with PD-L1 expression in tumor cells. The published literature on *UPP1* in liver cancer identifies it as a prognostic biomarker^27^ but, to our knowledge, offers no mechanistic account of why high *UPP1* predicts poor outcome. SMART’s results thus point to a possible mechanistic basis for this prognostic role.

### SMART supports discovery in a rare cancer

Because rare cancers have far fewer studies and less publicly available data^28^, findings from better-studied cancers can be a useful source of hypotheses for them. We therefore evaluated transfer from each of the 121 source studies to an scRNA-seq dataset of pleural mesothelioma, a rare, aggressive cancer with few treatment options^29^ (121 reports; Fig. 3d). We describe two of the results below.

A study of intestinal regeneration and colorectal tumorigenesis^30^ showed that *CYP27A1*-driven liver X receptor (LXR) activation restrains tumorigenesis through adaptive B and CD8 T cell responses. SMART found that *CYP27A1* expression correlated positively with the LXR target gene *ABCA1* and with the proportion of CD8 T cells in epithelioid tumors (Supplementary Data 11). *CYP27A1* has already been reported as a favorable prognostic marker in mesothelioma^31^, but, to our knowledge, why it carries this association has not been examined. In lung adenocarcinoma, higher *CYP27A1* expression has been linked to greater CD8 T cell infiltration^32^. The T cell association found by SMART may therefore point to a similar immune explanation for the prognostic value of *CYP27A1* in pleural mesothelioma.

ORACLE also transferred to pleural mesothelioma (Supplementary Data 11). In TCGA-MESO, high ORACLE scores were associated with worse survival (HR = 3.04, 95% CI 1.78–5.18, P = 2.2 × 10⁻⁵), consistent with a previous independent study^7^, and ORACLE was more strongly associated with survival than all 1,000 proliferation-matched random signatures (Supplementary Note 5 and Supplementary Fig. 2).

### SMART assembles tested mechanistic links into new hypotheses

Because SMART decomposes each source study’s mechanism into testable claims, the 726 reports yield a large collection of tested claims, each traceable to a specific paper. We refer to a claim as transferable when it is supported or partially supported. Claims involving the same gene can be grouped together, and claims that share a gene can be linked into a network. We illustrate both below in glioblastoma (Fig. 4).

**Figure 4.**
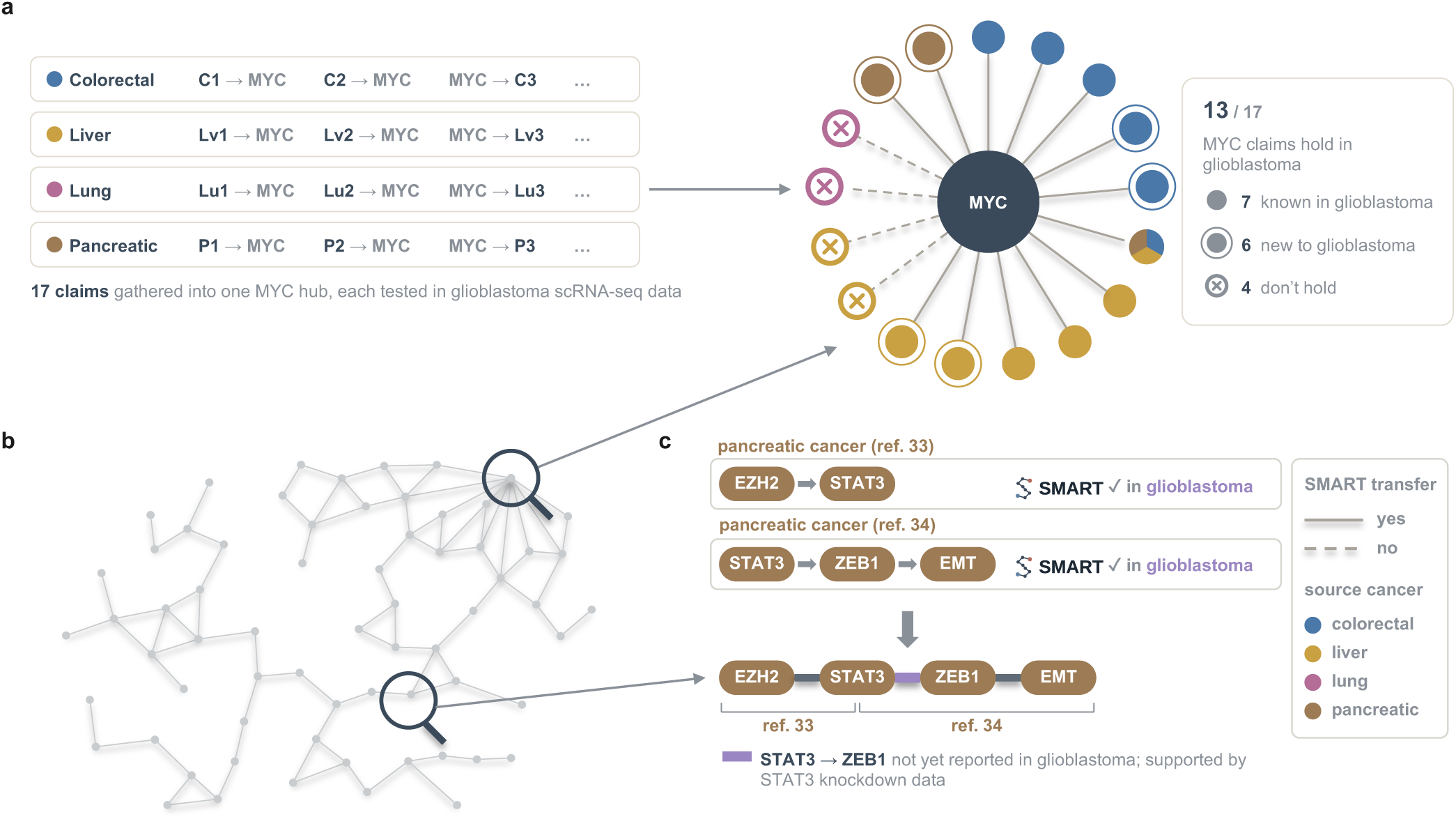
SMART links transferable claims across papers into new mechanistic hypotheses. Claims from the non-glioblastoma source studies were tested by SMART in the glioblastoma dataset and then regrouped by the genes, pathways and cell types they involve. **(a)** Claims involving *MYC* from non-glioblastoma source studies (left; labels are schematic, and the claims are listed in Supplementary Table 6), each tested by SMART in glioblastoma scRNA-seq data. Of 17 claims, 13 transfer (solid lines) and 4 do not (dashed lines, crossed circles). Among those that transfer, filled circles mark claims already reported in glioblastoma and double circles mark claims that are new. Circle color gives the cancer of the source study; the node divided into three colors marks a claim reported in colorectal, liver and pancreatic cancer studies. **(b)** Schematic of the network of transferable claims in glioblastoma; nodes are genes, pathways and cell types shared between claims and edges are claims. Arrows lead from the network to the enlarged views in panels a and c; the full network is available in CancerWiki. **(c)** An example path. Top, two claims from separate pancreatic cancer studies^33,34^, each transferring to glioblastoma (check marks), share *STAT3*. Bottom, the joined path *EZH2* → *STAT3* → *ZEB1* → EMT; brackets mark the source study of each segment. The purple *STAT3* → *ZEB1* link, not yet reported in glioblastoma, is supported by public *STAT3* knockdown data (Methods).

Across the non-glioblastoma source studies, 17 distinct mechanistic claims involve *MYC*, and SMART found that 13 of them transferred to glioblastoma (Fig. 4a and Supplementary Table 6). In our literature search, seven of these are already documented in glioblastoma. The remaining six have, to our knowledge, not been described in glioblastoma, making them promising leads for experimental follow-up.

Linking transferable claims that share a gene, pathway or cell type yields a mechanism network (216 claims from non-glioblastoma studies; Fig. 4b). A path through this network is a mechanism assembled from claims in separate papers. For example, *EZH2* → *STAT3* from a pancreatic cancer study^33^ and *STAT3* → *ZEB1* → EMT from a second pancreatic cancer study^34^, both transferable to glioblastoma, share *STAT3* and join into the path *EZH2* → *STAT3* → *ZEB1* → EMT (Fig. 4c). In glioblastoma, the *EZH2* → *STAT3*^35^ and *ZEB1* → EMT^36^ links are already established, and only *STAT3* → *ZEB1* is, to our knowledge, unreported, making it a candidate link between *EZH2* and EMT in glioblastoma. We therefore re-analyzed public data from a *STAT3* knockdown study in glioblastoma cell lines^37^ and found that knocking down *STAT3* reduced *ZEB1* expression (up to 1.7-fold, P ≤ 0.01; Methods). The full network is available in CancerWiki (Data availability).

### SMART suggests shared ovarian–pancreatic biology

The 726 reports can also be aggregated to ask which pairs of cancers exchange findings most often (Fig. 5a). Ovarian and pancreatic cancer stood out as the only pair where each was the top-ranked source of transferable findings for the other. Findings from ovarian studies transferred to pancreatic cancer in 8 of 12 cases (67%; other sources, 38–63%), and findings from pancreatic studies transferred to ovarian cancer in 9 of 15 cases (60%; other sources, 33–59%). Developmental origin does not readily explain this: ovarian studies transferred to pancreatic cancer more often than studies from liver, lung and colorectal cancer, which share the endodermal origin of pancreatic cancer^38^, whereas ovarian cancer does not^39^ (Fig. 5b).

**Figure 5.**
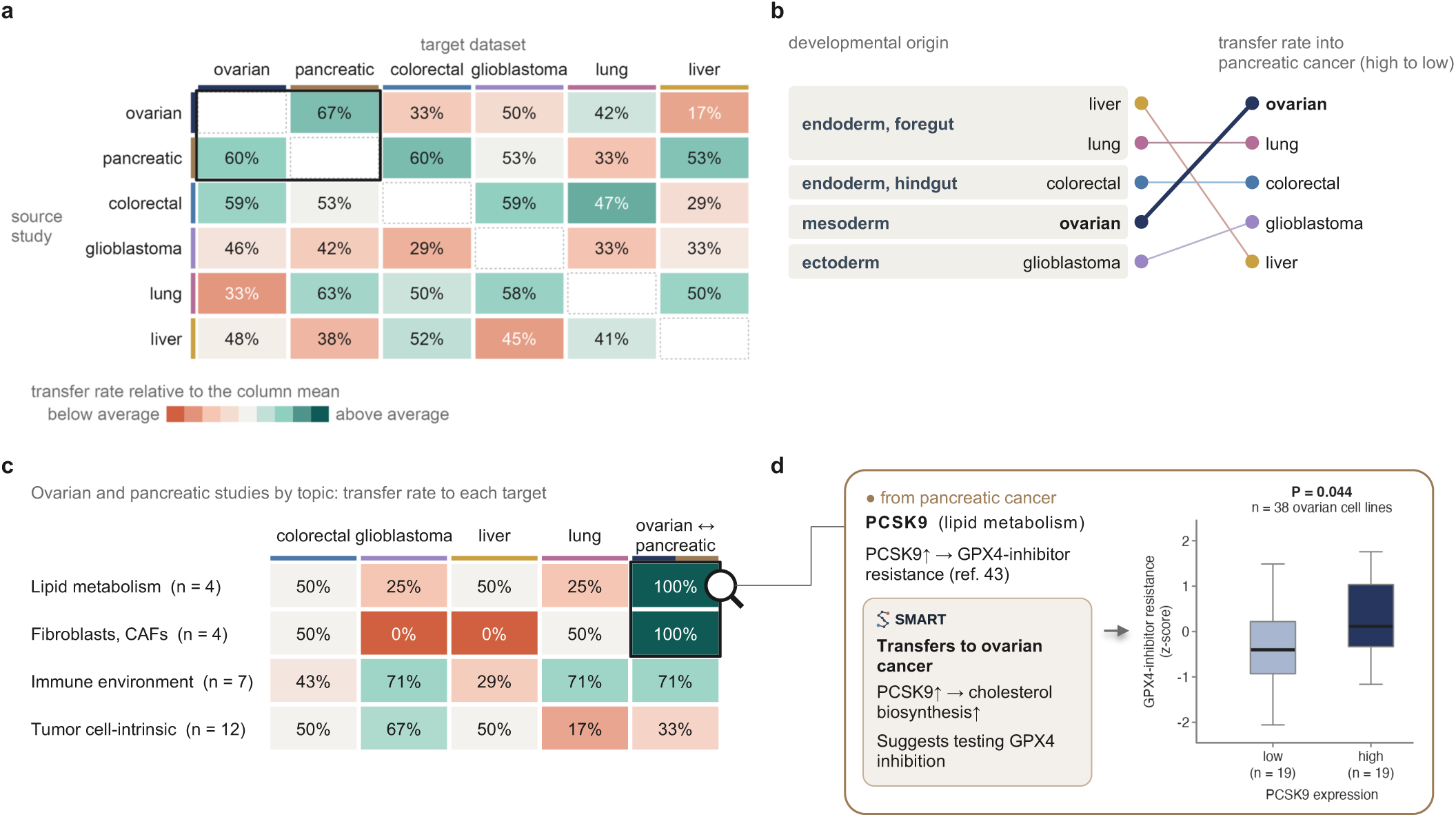
Aggregated transfer suggests shared ovarian–pancreatic biology. The 726 reports of Fig. 3a were aggregated by the cancer of the source study and of the target dataset to identify the pairs of cancers that exchange findings most often. **(a)** Transfer rates between cancers, by source cancer (rows) and target cancer (columns); a report was counted as a transfer when SMART assigned a transferability score of Middle-high or High, and Infeasible pairs were counted as non-transfers. Each cell gives the percentage of source studies that transferred and is colored relative to the column mean, from below to above average. The outlined cells mark ovarian and pancreatic cancer, each of which is the top-ranked source for the other; pairs in which source and target are the same cancer are not shown. Cancer colors are used throughout panels a–d. **(b)** Transfer into pancreatic cancer is not readily explained by developmental origin. Source cancers are ordered by developmental proximity to the pancreas (left) and ranked by transfer rate into pancreatic cancer (right). **(c)** Transfer rates of the 27 ovarian and pancreatic source studies, grouped by topic; n, number of studies. Cells are colored relative to the row mean, using the color scale of panel a. The first four columns give transfer into colorectal cancer, glioblastoma, liver cancer and lung cancer; the last gives transfer between ovarian and pancreatic cancer (ovarian studies into pancreatic cancer and pancreatic studies into ovarian cancer). Outlined cells mark topics in which all studies transferred between the two cancers; the magnifier points to panel d. **(d)** An example from the lipid metabolism group. A pancreatic cancer study^43^ reported that *PCSK9*-high cells are more resistant to GPX4 inhibitors than *PCSK9*-low cells, and SMART found that the cholesterol biosynthesis arm of the mechanism transferred to ovarian cancer. Right, resistance to GPX4 inhibitors (mean of z-scored areas under the dose–response curve for ML210, RSL3 and ML162) in ovarian cancer cell lines with *PCSK9* expression at or below the median (low, n = 19) or above it (high, n = 19). P, two-sided Wilcoxon rank-sum test. The full report is provided in Supplementary Data 12 and the analysis in Supplementary Note 6.

We therefore grouped the 27 ovarian and pancreatic studies by topic (Fig. 5c and Supplementary Table 7). All four studies of lipid metabolism and all four studies of fibroblasts transferred between ovarian and pancreatic cancer, whereas the same studies transferred to the other targets in at most half of cases. In contrast, only 4 of the 12 studies of tumor cell-intrinsic programs transferred between ovarian and pancreatic cancer, so the high transfer rate largely reflects the lipid metabolism and fibroblast findings. Both cancers grow in contact with adipose tissue, which supplies fatty acids to tumor cells^40,41^ and undergoes fibrotic remodeling as the tumor invades^40,42^. Ovarian and pancreatic cancer may therefore share lipid and stromal biology, and lipid- or stroma-targeted treatments developed for one may be worth testing in the other.

One example is *PCSK9*. A pancreatic cancer study^43^ reported that *PCSK9*-high cells synthesize cholesterol intermediates that protect against ferroptosis and are more resistant to GPX4 inhibitors. SMART found that in ovarian cancer, *PCSK9*-high tumor cells also showed higher cholesterol biosynthesis activity, and suggested testing GPX4 inhibition (Supplementary Data 12). The source study had reported an association between PCSK9 status and sensitivity to GPX4 inhibitors across cancer cell lines pooled from all lineages^43^; we further tested whether it holds within ovarian cancer. In drug-response data from 38 ovarian cancer cell lines, those with high *PCSK9* expression were more resistant to GPX4 inhibitors than those with low expression (P = 0.044), and higher *PCSK9* expression correlated with greater resistance (Spearman ρ = 0.39, P = 0.015; Fig. 5d, Supplementary Note 6 and Supplementary Fig. 3). Among all 407 compounds tested, the three GPX4 inhibitors ranked 19th, 27th and 42nd for this correlation (top 4.7%, 6.6% and 10.3%; Supplementary Note 6).

## Discussion

Published cancer studies generate numerous findings with potential relevance beyond their original cancer type, yet systematically evaluating these findings in other cancers requires substantial time, expertise, and computational resources. This makes discovery transfer particularly well suited to agentic automation. Here, we demonstrate that discovery transfer can be automated using SMART, a multi-agent large language model system that extracts and tests published findings in new cancer contexts. Manual audits confirmed the accuracy of 97% of evaluated outputs, and application to 121 studies across six cancer types identified transferable findings in 46% of cross-cancer reports.

Identifying findings that transfer across cancers can reveal shared biology and extend the relevance of published discoveries beyond their original setting. As shown here, SMART identified transferable prognostic signatures, conserved biological mechanisms, and potential therapeutic opportunities, demonstrating how discoveries made in one cancer can inform research in others.

Findings that do not transfer are also informative. A claim that lacks support in a new cancer is rarely reported, yet it tells investigators which mechanisms are unlikely to be worth pursuing there. For UPP1, SMART found no support for transfer to ovarian cancer, in agreement with a pan-cancer analysis^27^, and in glioblastoma it found that UPP1 was not linked to PI3K/AKT signaling, contrary to the source study, and suggested NF-κB signaling instead.

Discovery transfer may be particularly valuable for rare cancers, where limited research makes findings from more frequently studied cancers an important source of new hypotheses. In pleural mesothelioma, the prognostic value of ORACLE appeared in the literature more than four years after the signature was published^7^. SMART flagged the same transfer in a single pass over all 121 source studies. Notably, the pleural mesothelioma study found that ORACLE predicted survival slightly better than a signature developed specifically for pleural mesothelioma^7^, illustrating how findings from well-studied cancers can serve rare ones. SMART also linked CYP27A1, a previously reported prognostic marker in mesothelioma, to T cell infiltration in epithelioid tumors, suggesting a possible explanation for its prognostic association that, to our knowledge, had not been reported.

Evaluating transfer systematically, rather than through selective follow-up of a few studies, also allows results to be aggregated. Across 726 reports, the proportion of findings that transferred depended on the cancer pair and was not readily explained by developmental origin: ovarian and pancreatic cancer were each the top-ranked source for the other, and this was concentrated in findings on lipid metabolism and fibroblasts rather than on tumor cell-intrinsic programs.

Aggregation also operates at the level of individual claims. Because every claim is tested separately, claims from different papers can be regrouped by gene or joined through shared genes into a network, yielding hypotheses that no single source study states, such as an EZH2– STAT3–ZEB1–EMT axis in glioblastoma. Of its links, only STAT3–ZEB1 had not been reported in glioblastoma, and it was supported by public knockdown data.

Although scRNA-seq data cannot by themselves establish causal relationships, the associations that SMART tests are not chosen arbitrarily. They are derived from mechanistic claims in published studies, some of which are supported by functional experiments in their original biological context. SMART asks whether the molecular and cellular relationships expected under these mechanisms are also observed in a new context. Such evidence does not show that the original mechanism operates in the target setting, but it provides a biologically grounded basis for prioritizing specific hypotheses for functional validation and for designing targeted experiments.

Because each report runs independently, SMART can test many study–dataset pairs in parallel. A single report takes approximately 1.5 hours, and the 726 reports here were generated in 7 days. By comparison, even assuming an expert analyst needed only one day per pair, the same number of tests would take around two years.

In spite of SMART’s advantages, several limitations remain. Support in scRNA-seq data is not equivalent to functional validation, particularly for mechanisms that act through protein activity or spatial organization. Likewise, transfer in scRNA-seq data does not by itself establish prognostic value: ORACLE transferred to glioblastoma and colorectal cancer, but its survival association was not significant in those cohorts. Conversely, lack of support may reflect limited power, missing cell populations or incomplete metadata rather than true biological absence. The analyses used one target dataset for each of seven cancer types and source studies from Nature Portfolio journals (2024–2025), and some cancers were represented by few studies, so differences in transfer rate between cancer pairs should be interpreted with caution. As with any AI-based system, investigator review remains important, and SMART retains every intermediate output so that each step can be checked.

In summary, agentic discovery transfer makes it both possible and practical to test promising published cancer findings across new cancer contexts at scale. Each study can then not only inform its own cancer but also guide subsequent research investigations in every cancer to which its findings transfer. In this way, SMART increases the capacity of cancer researchers to systematically test published findings in their own data, with the ultimate goal of speeding the translation of a finding in one cancer into benefit across cancers.

## Methods

### Details on implementing SMART

SMART is implemented as a multi-agent workflow using Claude Code^15^, in which a shell script runs twenty specialized agents, organized into six modules, in a fixed order and stops if the Transferability Assessment judges the transfer infeasible. Each agent is defined by a structured prompt that specifies its role and the format of the file it must produce. Many prompts also carry a record of the errors that agents most commonly make, compiled through expert review of repeated runs, allowing each agent to check its own work against failure modes observed during development. Agents run sequentially within a module and communicate only through files, and the location and structure of those files are fixed in advance, so that every intermediate decision is retained on disk and can be inspected or re-run independently of the agent that produced it.

Each module reads the files written by the modules before it and writes its own output to disk in whichever format suits that stage, such as plain-text reports, JSON status files, CSV tables, R scripts, figures and rendered HTML reports. An overview of each module is given below, followed by details of the agents that make up each module.

### Results Extraction

SMART identifies the source study’s clinically relevant findings and mechanistic claims, verifying each against the paper text, the figure captions and the figure images, and then resolves every biological term the claims depend on into an scRNA-seq operational definition based on the source manuscript. The inputs are the manuscript text, the figure captions and images, the reference list and the supplementary files, which SMART retrieves from the article linked by the PubMed URL. The outputs are a paper summary and a definition table that maps each term to an scRNA-seq operational definition, such as a gene list or a named pathway. Users who have already summarized a study can supply these files directly, in which case the module is skipped.

### Data Inspection

SMART prepares and describes the target dataset before any analysis is planned. The input is the user-supplied dataset, and the output is a data inspection summary listing every metadata column with its type, distribution and representative values, the cell types present and expression-level summaries. Users who have already characterized their dataset can supply this summary directly, in which case the module is skipped.

### Transferability Assessment

SMART determines whether the source study’s mechanism can be tested at all in the target dataset. The inputs are the paper summary and the data inspection summary, and the output is a binary proceed decision with a one-sentence reason. When the decision is negative, the pipeline stops, so that computation is not spent on studies whose core logic the dataset cannot address.

### Task Plan

SMART translates the verified findings into a parameterized execution plan. The inputs are the paper summary, the definition table, the target dataset and the data inspection summary. The outputs are any additional cell-type annotations the definitions require, together with an execution plan consisting of the required preprocessing and analysis steps, each paired with a concrete function call whose column names and cell-type labels are taken from the target dataset and whose gene lists are taken from the definition table.

### Code and Analyze

SMART translates the planned analysis tasks into executable R scripts and runs them. The input is the execution plan, and the outputs are a preprocessing script, an analysis script, the resulting figures and statistical tables and a correction record documenting any issue discovered during the run together with the fix the module applied to make the script runnable. Scripts are assembled from a curated codebase of scRNA-seq analysis functions rather than written from scratch, which constrains agent behavior and makes results comparable across transfer tasks. Differential-expression log fold changes follow the alphabetical order of the group labels (Supplementary Information).

### Report

SMART interprets the computational output and compiles it into a structured report. The inputs are the execution plan, the figures, the statistical tables, the paper summary and the data inspection summary, and the output is a rendered HTML report that assesses each claim as supported, partially supported or not supported, assigns an overall transferability score, flags new and contradictory signals and recommends follow-up analyses or experiments.

The agents constituting each module are described below.

**Results Extraction** consists of nine agents in three stages: drafting, verification and definition resolution.

### Summary Draft agent

reads the manuscript with the single goal of understanding what the study found, and writes a structured draft stating the clinically relevant findings and the central causal chain. The agent does not inspect figures or extract statistics; its task is to capture the gist of the study, and detailed verification is left to the agents that follow. It also records every figure-referencing sentence in the paper in a mapping file that the verification agents can query.

### Clinical Findings agent

verifies the clinically relevant findings drafted upstream. Each of these findings states a patient-level or translational result, such as an expression difference between diseased and normal tissue, a response to a therapeutic agent or an association with survival or outcome, and carries the subfigure-level citations that support it. The agent therefore checks both the citations and the wording of each finding: it labels each one by comparing its citations against the mapping file, reads the surrounding paper text for any finding flagged as uncertain and corrects missing references, incorrect subfigure ranges, overstated modifiers and overclaimed scope.

### Mechanistic Claims agent

decomposes the mechanism section into individual causal claims, locates each one in the manuscript by keyword and figure-reference search and evaluates it along six dimensions: causal direction, meaning whether the draft has A acting on B or the reverse; claim certainty, meaning whether the draft matches the confidence the paper itself expresses; feedback loops; pathway accuracy; completeness of parallel mechanisms; and composite inference, meaning whether combining separately supported findings has produced a claim the paper never makes. Claims that cannot be located are searched for by reading the relevant Results or Discussion section in full before they are corrected or removed.

### Mechanism Figure agent

checks whether the authors summarized their own mechanism in a schematic figure, and if so uses that figure to verify the mechanistic claims. The agent screens the figure captions for a candidate and confirms it by viewing the image, then extracts every element the figure shows and compares the mechanism section against it, adding elements the draft omitted, correcting contradictions and removing or downgrading content that neither the figure nor the text supports.

### Figure Reference agent

audits the figure references attached to each clinically relevant finding, reading the figure images themselves to confirm that each cited panel shows what the finding claims. Two problems motivate this step. Papers often cite a group of panels for a group of findings, so when the summary keeps only part of that group, the panels that support the other findings have to be removed. Authors also occasionally misassign a reference, in which case the caption alone is not sufficient and the correct panel can be identified only by inspecting the image.

### Definition Extraction agent

enumerates every biological term the paper summary depends on and decides which of them require an explicit definition before the findings can be tested in another dataset. Terms with no transcriptional readout are marked as not applicable, and single named genes are treated as unambiguous. The agent collects the remaining terms, such as cell subtypes, gene sets, pathways, signatures and phenotypes, into a list of terms that need a detailed check, which the agents below then resolve.

### Definition Lookup agent

resolves each flagged term into a single usable definition by searching the main text and the methods, and then opening supplementary tables, figures and documents where the paper points to them. A fixed reliability hierarchy selects among competing candidates; for example, gene sets validated by follow-up experiments take priority over the computational enrichment results that produced them. Aliases are converted to official gene symbols. Terms the agent cannot resolve by searching the source study alone are routed to one of the two agents below.

### External Reference agent

resolves terms whose definitions are given in a cited paper rather than in the source study or its supplementary files. The agent locates the referenced publication and extracts the term definition from its text or supplementary files.

### External Package agent

resolves terms that the source study defined through a specialized package. Such packages are typically released alongside a paper that proposes its own classifier for a set of cell subtypes. The agent installs the package, determines how the study applied it and writes a tested usage guide so that the same classifier or scoring method can be applied to the target dataset.

**Data Inspection** consists of two agents.

### Preprocess agent

prepares the target dataset for analysis. When the dataset is supplied as an h5ad file, the agent converts it to a Seurat object, retaining the raw counts and all cell metadata. When unprocessed counts are supplied together with the user’s metadata, the agent assembles them into a Seurat object, normalizes the counts and uses the supplied metadata as given; when no metadata are supplied, it also performs quality control, clustering and cell-type annotation. When a processed object is supplied, the agent leaves the expression data and existing annotations unchanged and makes only the edits that downstream analysis requires. For example, cells with a missing label in the finest cell-type annotation are assigned their label from a coarser annotation, and a patient identifier column is added when patient identity is encoded in the cell barcodes but absent from the metadata.

### Data Inspection agent

loads the prepared dataset and produces the data inspection summary. The agent describes only what is present, including every metadata column with its type, distribution and sample values, the cell-type composition and expression summaries. It performs no filtering, transformation or interpretation, so that downstream decisions are made from a complete view of the data.

**Transferability Assessment** consists of a single agent that checks that the cell types the source mechanism requires have counterparts in the dataset, and that the mechanism has a transcriptional component that scRNA-seq can observe. The agent performs a gate check rather than a scientific assessment; whether the mechanism actually transfers is determined by the downstream modules.

**Task Plan** consists of three agents.

### Data Process agent

subclusters broad cell populations to produce the finer subtypes the definitions require. A dataset may label its cells only as T cells and macrophages, for example, while testing the paper requires regulatory T cells and SPP1⁺ macrophages. The agent compares the definition table against the data inspection summary and resolves each missing subtype using the definition the paper supplied for it. New annotation columns are written to a separate table, and the original dataset is not modified.

### Task Brainstorm agent

enumerates the analyses needed to test each finding in two stages. The first stage asks what the ideal test would be, given a dataset that contained everything the paper’s findings call for, and the agent is therefore not allowed to read the data inspection summary at this point. It decomposes each paper finding into standard analysis patterns, so that, for example, a claim that gene A activates gene B becomes a correlation between A and B across samples, together with a comparison of B between the samples with high and low A. Claims that operate at the protein or metabolite level are given an mRNA proxy where one is reasonable, with an explicit note that the proxy may not yield a result. The second stage reads the data inspection summary and revises this ideal plan against the dataset in hand, sorting each analysis into directly testable, testable after preprocessing or not testable, and dropping what the dataset cannot support. It then adds stratifications that the available metadata make biologically meaningful, such as repeating an analysis separately by sex or by tumor grade, even when the source study did not report on these conditions. Finally, it proposes a small number of additional analyses that the dataset enables beyond the paper’s claims.

### Task Revise agent

converts the categorized plan into a unified execution plan. The agent first identifies every preprocessing step the analyses require and merges duplicates so that each score or sample grouping is computed once, recording which analyses depend on it, and then decomposes each analysis into concrete function calls. Every parameter is filled in with a real value read from the target dataset, using its own column names, cell-type labels and condition values rather than placeholders. The resulting plan leaves the Code Writing agent no remaining decisions about which functions to call or what parameters to pass.

**Code and Analyze** consists of two agents operating against a shared analysis codebase.

### Code Writing agent

translates the execution plan into two executable scripts. The preprocessing script computes the variables the analyses are defined over, such as pathway and signature scores and the split of samples into high and low groups for a chosen feature, and it also merges in the subtype labels the Data Process agent produced earlier. The distinction is that the Data Process agent establishes cell identity, whereas the preprocessing script derives the quantities each analysis compares, such as a pathway activity score. Rather than saving a new copy of the dataset, the preprocessing script writes only the updated metadata to a table, and the analysis script loads the original dataset, joins that table onto it and runs every analysis task in turn. The Code Writing agent builds both scripts from a curated single-cell analysis codebase rather than writing functions from scratch. Each function is implemented in R and is paired with usage documentation that the agent reads before generating any code. The codebase provides functions for dataset inspection, module score computation, sample stratification by feature, gene-set retrieval from ontology and signature databases, per-sample feature correlation, group comparison, differential expression, cell composition comparison and enrichment analysis. Restricting the agent to these predefined functions improves accuracy, efficiency and consistency^44^, and it also simplifies validation, because verifying an analysis reduces to checking that the correct function was called with appropriate parameters. Function implementations and documentation are provided in the SMART repository.

### Code Execution agent

runs both scripts, captures their logs and debugs whatever fails, patching the script and re-running until the analysis completes. Patches are written to separate corrected files so that the original scripts remain intact. The number of correction attempts is capped, and every patch is recorded with the error that triggered it and the outcome of the re-run.

**Report** consists of three agents.

### Report Brainstorm agent

walks through every analysis step in order, locates its output and interprets it against the source mechanism, and then synthesizes the per-step observations into an overall assessment. The agent assigns one of four transferability scores (see Transferability score).

### Final Report agent

decomposes the paper summary into granular claims, each small enough for a clear yes, no or partial verdict on whether it transfers, maps each claim to its supporting statistics and to one or two selected figures and generates a seven-section report covering the study and dataset overview, a per-claim validation table, per-claim transfer results with figures, contradictions and new findings, interpretation of those contradictions, the transferability assessment score and recommendations that follow from that score. A fixed theme and structure keep reports comparable across studies and datasets. Reports are rendered with R Markdown.

### Report Revision agent

checks the generated report against its own sources, verifying that every reported number matches the underlying statistical output, that each figure supports the claim it illustrates, that no claim from the assessment was dropped and that the interpretation does not exceed what the results establish. The agent returns a revised report together with a log of every change made.

SMART calls each agent as the pipeline progresses and writes its output to disk, so that a completed run retains the summary drafts, the verification reports, the definition table, the execution plan, the generated scripts, the correction record and the per-step interpretations that underlie the final report. Full agent prompts, reference files and the complete pipeline specification are available in the SMART repository (https://github.com/cancer-wiki/SMART).

### Transferability score

SMART assigns each source study–target dataset pair one of four transferability scores, graded by how much of the study’s mechanistic chain holds in the target dataset. High means the core mechanistic chain transfers broadly, with strong statistical support. Middle-high means the key downstream claims transfer, although parts of the chain do not hold, hold only under specific conditions or follow an alternative route. Middle-low means some downstream signal appears only under specific conditions, while the core mechanism does not hold broadly. Low means only the canonical upstream wiring is preserved, while the downstream claims that constitute the study’s new contribution fail. Pairs for which the Transferability Assessment module determined that the source mechanism could not be tested in the target dataset were labeled Infeasible and not scored. For aggregate analyses, a pair was counted as a transfer when the score was Middle-high or High; pairs scored Middle-low or Low, and pairs labeled Infeasible, were counted as non-transfers. At the level of individual claims, SMART labels each claim as supported, partially supported or not supported; we refer to a claim as transferable when it is supported or partially supported.

### Target datasets and preprocessing

Seven public scRNA-seq datasets were used as targets, one per cancer type, together with the dataset of Qi et al.^22^ for the positive control (sources in Data availability). Each dataset was used with the cell-type annotations and metadata provided by its source. Because of memory constraints, the ovarian cancer dataset was restricted to 11 of its 34 patients, selected at random. This study used only publicly available, de-identified data, and no ethics approval was required.

### Comparison with general-purpose agents and manual validation

The comparator systems, prompts, evaluation tasks and scoring used to compare SMART with general-purpose large language models (LLMs) and LLM-based agents are described in Supplementary Notes 1 and 2, Supplementary Table 1, and Supplementary Data 6, and the outputs of each system are provided in Supplementary Data 1–5.

For each manually validated case (Supplementary Note 4), we audited SMART’s outputs against the source manuscript and the target dataset. For the Results Extraction module, each clinically relevant finding and mechanistic claim returned by SMART was paired with the original text from the source paper to determine whether the extraction was accurate. Biological term definitions produced by the same module were evaluated similarly, with each definition checked against the relevant manuscript text, figure or supplementary file. For the Data Inspection module, we confirmed that the reported cell types and metadata matched the target dataset. For the Transferability Assessment module, we recorded whether SMART’s decision to proceed agreed with expert assessment. For each testable claim, we determined whether SMART executed a proper test and stated its result in the final report. The 10 cases, the transferability score assigned to each and the comparison across three independent runs are listed in Supplementary Tables 2 and 3.

To construct decoys, we used the six source studies behind the 15 reports featured in this study (Qi et al., ORACLE, UPP1, CYP27A1, PCSK9 and hnRNPL; Supplementary Table 4): ORACLE against six targets, UPP1 against five, and each of the other four against one target. For each, we built a decoy paper summary by reversing the direction of every directional statement in the clinically relevant findings and mechanistic claims; for example, “correlates with” became “is inversely correlated with”, and “enriched” became “depleted”. The mechanism section was rewritten as explicit statements of the reversed associations, and each decoy was checked for internal consistency so that no chain of reversed steps implied the original direction. Decoy summaries were run through SMART from the Task Plan module onward, using the original definition tables and data inspection summaries, against the same target datasets as the original 15 reports (Supplementary Table 4).

### Large-scale application and CancerWiki network construction

Candidate source studies were identified in October 2025 by a keyword search of titles and abstracts in PubMed, restricted to articles published in 2024–2025 in Nature, Nature Cancer, Nature Communications, Nature Genetics, Nature Immunology or Nature Medicine, for scRNA-seq studies of colorectal cancer, glioblastoma, liver cancer, lung cancer, ovarian cancer or pancreatic cancer. Each candidate was reviewed manually and included if (1) its major result was a biological discovery about the cancer type of interest and (2) that discovery was at least partially testable in scRNA-seq data; 121 studies met both criteria (Supplementary Table 5). Selection was based on the publications alone and was completed before SMART was applied. The transfer rates reported here refer only to studies with findings testable in scRNA-seq data. Each source study was run against each of the six target datasets (726 reports) and against the mesothelioma dataset (121 reports). The transfer rate for a source cancer and a target cancer was the proportion of source studies counted as transfers, computed for pairs of different cancers. The ovarian and pancreatic source studies were each assigned to one topic (lipid metabolism, fibroblasts, immune environment or tumor cell-intrinsic programs) by manual reading of the study (Supplementary Table 7). Cancers were grouped by the germ layer from which the tissue of origin derives: endoderm for colorectal, liver, lung and pancreatic cancer^38^, mesoderm for ovarian cancer^39^ and ectoderm for glioblastoma. For the mechanism network, the transferable claims (supported or partially supported) of each non-glioblastoma source study evaluated in the glioblastoma dataset were extracted, and claims from different studies were joined at the genes, pathways and cell types they share, so that these form nodes and claims form edges. We write A → B for a claim that A acts on B.

### Survival analyses in TCGA cohorts

Bulk RNA-sequencing data and overall survival for the TCGA cohorts were obtained from the UCSC Xena TCGA hub^45^. ORACLE was applied as in the original publications^5,6^, without refitting or rescaling: for each patient’s primary tumor, the score was computed as the sum of the expression of the 23 signature genes (RSEM normalized counts, log2(x + 1)) weighted by the published coefficients. All 23 genes were present in every cohort. Patients were divided at the within-cohort median score. Following the original publications, analyses were restricted to stage I–III disease, except in TCGA-GBM, which is uniformly grade IV and was analyzed whole. TCGA-COAD and TCGA-READ were pooled as a single colorectal cancer cohort. Overall survival was the endpoint. Kaplan–Meier curves were compared with a two-sided log-rank test, and hazard ratios (high versus low) were estimated by univariable Cox proportional hazards regression, using the survival package (v3.5.7). Cohort sizes and full statistics are given in Supplementary Note 5 and Supplementary Fig. 2. To test whether proliferation explained these associations, ORACLE was compared with 1,000 random 23-gene signatures per cohort whose correlation with a Hallmark G2M proliferation score was within ±0.05 of ORACLE’s (Supplementary Note 5).

### Drug-sensitivity analysis

Drug-response data were taken from the Cancer Therapeutics Response Portal v2.1 (CTRP)^46,47^, downloaded from the NCI CTD² data portal (dataset CTRPv2.1_2016_pub_NatChemBiol_12_109). For three GPX4 inhibitors (ML210, RSL3 and ML162), the CTRP area under the dose–response curve was z-scored across ovarian cancer cell lines and averaged into a resistance score for each line. *PCSK9* expression was obtained from DepMap^48,49^ (DepMap Public 26Q1). The 38 ovarian cancer cell lines with both measurements were divided at the median *PCSK9* expression, and the two groups were compared by a two-sided Wilcoxon rank-sum test. Spearman’s rank correlation between *PCSK9* expression and the resistance score was also computed across all 38 lines. To assess specificity, the Spearman correlation between *PCSK9* expression and the area under the dose–response curve was computed for each of the 407 compounds tested in at least 20 of these lines, and compounds were ranked by correlation. Analyses used R 4.3.2; full details are given in Supplementary Note 6.

### *STAT3* knockdown data in glioblastoma

We re-analyzed public microarray data from *STAT3* knockdown with three independent shRNA hairpins in patient-derived glioblastoma lines^37^ (GEO accession GSE117905). Significance was assessed by a two-sided t-test. All four *ZEB1* probe sets decreased with each of the three hairpins (by 0.30–0.77 log2 units; P = 0.0004–0.010).

### Literature search

To assess whether a finding or claim had been reported in a target cancer, we searched Google Scholar using the gene or signature name together with the cancer type and manually reviewed the most relevant results. A finding was described as not previously reported when this search identified no study reporting it in the target cancer. These searches, performed in October 2026, were not systematic reviews, and such statements are accordingly qualified as being to our knowledge. This procedure was applied to ORACLE, *UPP1* and *CYP27A1* in the relevant cancers, to the *MYC* claims tested in glioblastoma, and to each link of the *EZH2*–*STAT3*–*ZEB1*– EMT path. The *MYC* claims, with the supporting publication for those already documented in glioblastoma, are listed in Supplementary Table 6; supporting publications for the other findings are cited in the text. For ORACLE, all studies citing the two ORACLE papers were retrieved from Google Scholar in October 2026, and each was reviewed manually for direct evaluation of the signature in another cancer type.

### Statistics and reproducibility

The Wilcoxon rank-sum test was used for the drug-sensitivity analysis, together with Spearman’s rank correlation between *PCSK9* expression and resistance, and the log-rank test and univariable Cox proportional hazards regression for survival analyses, a two-sided t-test for the *STAT3* knockdown analysis, and an empirical comparison with 1,000 proliferation-matched random signatures for ORACLE (Supplementary Note 5). Statistics reported from SMART’s analyses were computed with the functions of its curated codebase. No statistical method was used to predetermine sample size; sample sizes were set by the available source studies, public datasets and cell lines. Apart from the random selection of 11 of 34 patients in the ovarian cancer dataset because of memory constraints, no data were excluded. Randomization and blinding were not applicable. All agents used Claude Opus 4.6 with default settings. Each source study–target dataset pair in the large-scale application was run once; run-to-run variability was assessed separately by running each of the 10 manually validated cases three times (Supplementary Note 4 and Supplementary Table 3).

### Reporting summary

Further information on research design is available in the Nature Portfolio Reporting Summary linked to this article.

## Data availability

All datasets analyzed in this study are publicly available and were obtained from their original publications. The seven target single-cell RNA-sequencing datasets are available as follows: colorectal cancer, Gene Expression Omnibus (GEO) accession GSE132465; glioblastoma, processed object obtained from the Single Cell Portal (accession SCP1985); liver cancer, GEO accession GSE149614; lung cancer, GEO accession GSE127465; ovarian cancer, GEO accession GSE222556; pancreatic cancer, processed object obtained from Zenodo^50^ (https://zenodo.org/records/3969339); and pleural mesothelioma, GEO accession GSE327167.

The colorectal cancer dataset used for the positive-control analysis was obtained from the Supplementary Information of ref. 22.

Bulk RNA-sequencing and clinical data, including overall survival, for the TCGA-LIHC, TCGA-PAAD, TCGA-GBM, TCGA-OV, TCGA-COAD, TCGA-READ and TCGA-MESO cohorts were obtained from the UCSC Xena TCGA hub^45^ (https://tcga.xenahubs.net). Drug-response data (CTRP v2.1)^46,47^ were obtained from the NCI CTD² data portal (dataset CTRPv2.1_2016_pub_NatChemBiol_12_109) and cell-line expression data from DepMap^48,49^ (DepMap Public 26Q1; https://depmap.org/portal). Microarray data from *STAT3* knockdown in glioblastoma lines were obtained from GEO (accession GSE117905). The 121 source studies, with their DOIs and the transferability scores for all 726 reports, are listed in Supplementary Table 5.

The SMART reports and outputs discussed in this study are provided as Supplementary Data 5 and 8–12. The glioblastoma mechanism network is available in CancerWiki (https://cancer-wiki.org).

## Code availability

The SMART code, agent prompts and curated analysis codebase are available at https://github.com/cancer-wiki/SMART.

## Acknowledgements

This work was supported by the National Institute of General Medical Sciences of the National Institutes of Health (R35GM162239) and the National Institute of Allergy and Infectious Diseases of the National Institutes of Health (R21AI190872). The results shown here are in part based upon data generated by the TCGA Research Network: https://www.cancer.gov/tcga.

## Author contributions

L.C. and J.S. conceived the project. L.C. led the project, designed and implemented SMART, performed testing and interpreted the results. J.S. contributed to the biological interpretation of the results. A.G. contributed to method development. M.M.S. provided input on the biological background. Y.C., E.X. and P.K. curated the datasets. C.K. supervised the study. L.C., J.S. and C.K. wrote the manuscript. M.M.S. and M.D. reviewed the manuscript.

