## Supplementary Information for "Agentic Discovery Transfer Enables Cross-Cancer Discovery at Scale"

### Contents

### Supplementary Notes


| | | Page |
|---|---|---|
| **Note 1** | Illustrative example: why a structured workflow matters for discovery transfer | 3 |
| **Note 2** | Comparison of SMART's Results Extraction module with a general-purpose LLM | 8 |
| **Note 3** | Example output of SMART's Data Inspection module | 9 |
| **Note 4** | Manual validation of SMART in 10 cases | 12 |
| **Note 5** | ORACLE survival association in bulk cohorts | 15 |
| **Note 6** | *PCSK9* expression and GPX4-inhibitor sensitivity in ovarian cancer cell lines | 17 |


### Supplementary Tables

*Supplementary Tables 5 to 7 are provided as separate Excel files.*


| | | Page |
|---|---|---|
| **Table 1** | Definitions used by the four agents and SMART for the source study's key terms (in Supplementary Note 1) | 7 |
| **Table 2** | The 10 manually validated cases (in Supplementary Note 4) | 12 |
| **Table 3** | Transferability scores across three independent runs of the 10 validated cases (in Supplementary Note 4) | 13 |
| **Table 4** | Transferability scores of the original and decoy reports (in Supplementary Note 4) | 14 |
| **Table 5** | Source studies and transferability scores for all 726 SMART reports (Excel file) | 19 |
| **Table 6** | Mechanistic claims involving *MYC* and their transfer to glioblastoma (Excel file) | 19 |
| **Table 7** | Topic assignment of the ovarian and pancreatic source studies (Excel file) | 19 |

#### Supplementary Figures


| | | Page |
|---|---|---|
| **Fig. 1** | Definitions and claim coverage across the four agents and SMART (in Supplementary Note 1) | 6 |
| **Fig. 2** | ORACLE score and overall survival in six TCGA cohorts (in Supplementary Note 5) | 16 |
| **Fig. 3** | Correlation between *PCSK9* expression and GPX4-inhibitor resistance across 38 ovarian cancer cell lines (in Supplementary Note 6) | 18 |


#### Supplementary Data

*All Supplementary Data files are provided separately.*


| | | Page |
|---|---|---|
| **Data 1–4** | Outputs of the comparator agents | 20 |
| **Data 5** | SMART report for the example in Supplementary Note 1 | 20 |
| **Data 6** | Comparison of SMART's Results Extraction module with a general-purpose LLM | 20 |
| **Data 7** | Manual audit of the SMART workflow in the 10 validated cases | 20 |
| **Data 8** | SMART report for the positive control | 20 |
| **Data 9** | SMART reports on the transfer of ORACLE | 20 |
| **Data 10** | SMART reports on the transfer of UPP1 | 20 |
| **Data 11** | SMART reports on the transfer of CYP27A1 and ORACLE to pleural mesothelioma | 21 |
| **Data 12** | SMART report on the transfer of PCSK9 | 21 |

### Supplementary Notes

#### Supplementary Note 1. Illustrative example: why a structured workflow matters for discovery transfer

This example is not a benchmark of general-purpose agents, which are built for a different and broader purpose. It illustrates two failure modes, definition substitution and incomplete coverage of the paper's findings, that a task-specific workflow is designed to prevent.

Several recent LLM-based agents can execute biological analyses autonomously by combining large language model (LLM) reasoning with tool use and code execution. Biomni is a general-purpose biomedical AI agent that answers questions and executes analytical tasks across many subfields, from genetics and genomics to single-cell analysis[14]. Claude Code (Anthropic) is a general agentic coding assistant that calls tools and runs commands from user instructions[15]. Claude Science (Anthropic) is an AI workbench for scientific research in which an agent works across databases, code tools and compute on the user's own infrastructure[16]. Kosmos (Edison Scientific) is an autonomous AI scientist that, given a research objective, iterates between literature search, data analysis and hypothesis generation to produce a cited report[17].

Unlike these generalists, SMART is an agentic workflow designed for one task that recurs constantly in research practice, which we define as discovery transfer (determining whether results from one study transfer to a different dataset). Because SMART is defined as a workflow rather than a model, it could in principle be built on other LLMs; here we implement it with Claude Code running Opus 4.6.

##### Comparison setup

We gave the same discovery-transfer task to Biomni, Claude Code, Claude Science and Kosmos, and compared their output with that of SMART. The source study was Qin et al., which reports that hnRNPL phase separation activates *PIK3CB* transcription, promoting glycolysis and thereby tumor progression in ovarian cancer[51], and the target was the glioblastoma single-cell dataset used throughout this work (Single Cell Portal accession SCP1985). All agents received the same two inputs, the article URL and the target dataset, and the same single prompt, "Using this paper (https://www.nature.com/articles/s41467-025-60115-7), how do the clinically relevant findings and the mechanistic claims transfer to my attached data?", with no follow-up other than approving permission requests when prompted.

Claude Code and Claude Science were run with the same model as SMART (Claude Opus 4.6), so the difference in output reflects the workflow rather than the underlying model. Biomni was run in its Max mode, and Kosmos as the legacy Kosmos agent on the Edison Scientific platform.

The output of the four agents and SMART was compared against the source manuscript: whether a claim appears in the paper, whether a gene list matches the paper's, and whether each testable link in the paper was analyzed or missed.

##### Comparison results

Neither SMART nor any of the four general-purpose agents reported that the paper's mechanism transfers in full. Read alone, each of the general-purpose agents' reports is plausible. Checked against the paper, the four general-purpose agents differ in which claims they examine and how they define and test them, so a reader has no basis for trusting one over another. Two issues are common to all four.

*Definition substitution.* The source study validates 14 PI3K-AKT pathway genes as hnRNPL targets by RT-qPCR (Fig. 4i and Supplementary Fig. 5c of the source study). SMART used exactly these 14. In place of the paper's hnRNPL-regulated PI3K-AKT genes, Biomni, Claude Code and Claude Science each assembled their own list (12, 10 and 17 genes), and Kosmos used the 105-gene Hallmark PI3K-AKT-mTOR set. These match 5, 4, 4 and 1 of the paper's 14 genes respectively (Supplementary Fig. 1a, Supplementary Table 1). Kosmos states that the paper did not supply a transcript signature for PI3K-AKT, which overlooks these 14 genes. Biomni states that its PI3K-AKT gene set was curated from the paper text, although only 5 of its 12 genes are among the paper's 14 validated targets. Biomni then reports a positive association between *HNRNPL* and its PI3K-AKT list (correlation 0.68) as evidence that hnRNPL's regulation of the PI3K-AKT pathway transfers, and attributes it to *CDK6*, *MTOR*, *AKT3* and *PIK3R1*. Of these, only *CDK6* is among the paper's 14 genes. The association therefore rests mostly on genes the paper does not implicate.

*Coverage of the paper's own findings.* The paper functionally validates that hnRNPL promotes tumor progression, through proliferation and through migration and invasion. None of the four general-purpose agents extracted proliferation or migration and invasion as claims of the paper. Claude Code and Kosmos describe the endpoint only in general terms ("tumor progression", "ovarian cancer phenotypes"); Biomni raises proliferation only as future work, Claude Code only as a possible confounder of its own correlations, and Claude Science proposes "hnRNPL → *CDK6* → cell cycle progression" without testing it. None tested whether hnRNPL's effect on proliferation or on migration and invasion transfers. SMART extracted both phenotypes as claims and tested each. For proliferation, in primary glioblastoma, *HNRNPL* expression correlates with the fraction of glioma cells in S or G2/M phase, and genes upregulated with high *HNRNPL* expression are enriched for DNA replication, mitotic cell cycle phase transition and G1/S transition, with no glycolysis terms among them (Supplementary Data 5). SMART therefore proposed that in glioblastoma the hnRNPL–PIK3CB–AKT axis is linked to proliferation rather than to the glycolysis reported in ovarian cancer. Because only SMART tested proliferation, its result could be connected to earlier work on HNRNPH1, a related hnRNP protein that drives glioblastoma progression by regulating the splicing of cell cycle genes[52].

Supplementary Fig. 1b records which of the paper's claims the four agents and SMART extracted, and the definition each used. Of the seven claims shown, SMART extracted all seven; the four general-purpose agents extracted 3 to 5. Supplementary Table 1 gives each definition in detail. The full output of each agent is provided as Supplementary Data 1 to 4; the complete SMART report is provided as Supplementary Data 5. SMART's full workflow, from result extraction to report, was verified step by step (Supplementary Data 7).

#### Why accuracy matters for unsupervised use

An autonomous tool is useful only if its output need not be redone by hand. A reader can check a summary against the paper but not the analyses built on it, so a single visible error casts doubt on the rest. Because such errors are unpredictable, finding one does not reveal where the others are, and the safe response is to re-check everything, which removes the point of automating the task.

SMART therefore uses the paper's own claims and definitions, labels any proxy and tests every extracted claim, so that each step can be checked against the source study.

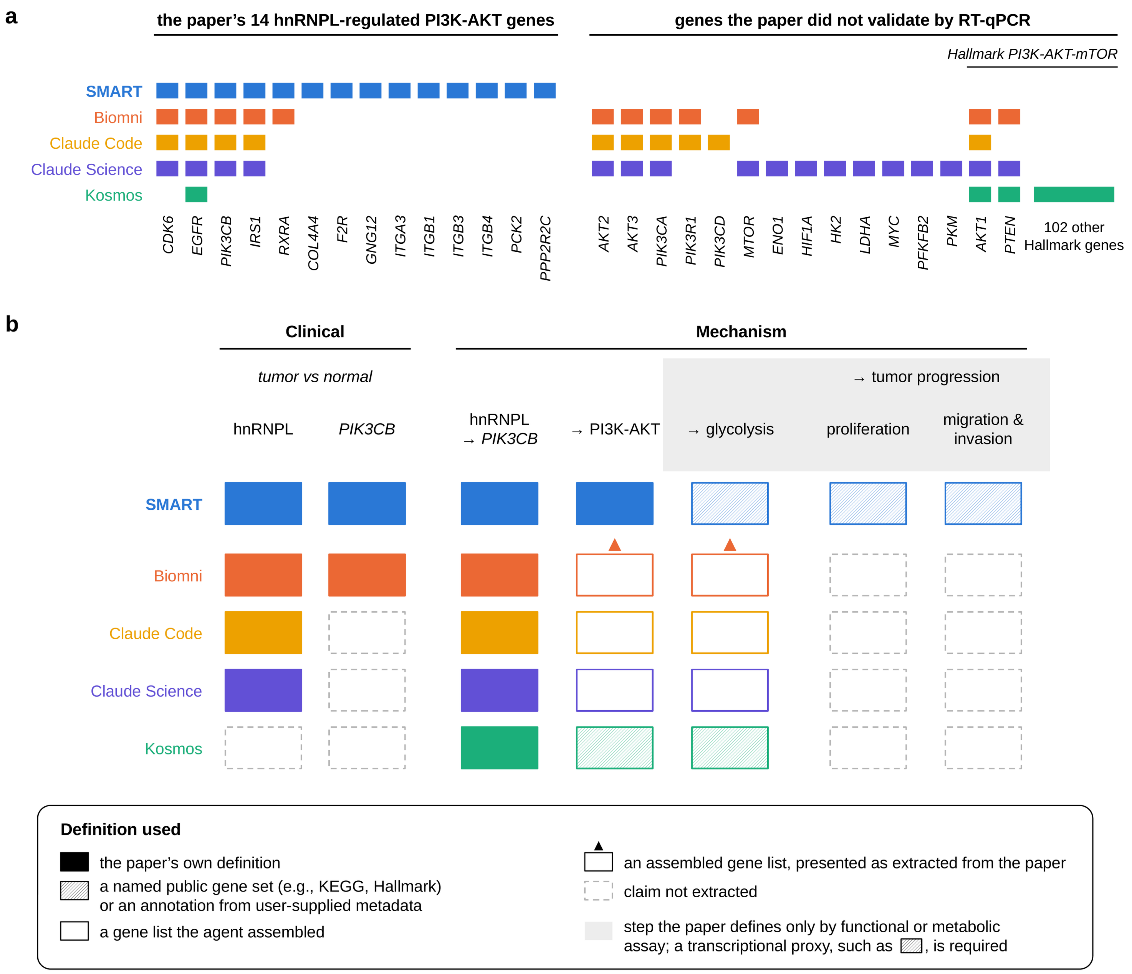


**Supplementary Fig. 1. Definitions and claim coverage across the four agents and SMART.** **a**, Gene sets used by SMART and the four agents for the paper's hnRNPL-regulated PI3K-AKT genes. The paper validates 14 PI3K-AKT pathway genes as hnRNPL targets by RT-qPCR; a filled cell means the gene was used. SMART used all 14 and nothing else. Biomni, Claude Code and Claude Science assembled their own lists around canonical PI3K-AKT genes; Kosmos used the 105-gene Hallmark PI3K-AKT-mTOR set, which shares one gene with the paper's 14 (*EGFR*). None of the four agents used nine of the paper's 14 genes. **b**, Claims of the source study extracted by each agent, and the definition used to test each. Columns follow the paper's chain from hnRNPL to tumor progression. Cell fill records where the definition came from: solid, the paper's own definition; hatched, a named public gene set such as KEGG or Hallmark, or an annotation from the user-supplied metadata of the target dataset; hollow, a gene list the agent assembled; hollow with a triangle, a gene list the agent assembled and presented as extracted from the paper; dashed outline, claim not extracted. Grey column headers mark steps the paper defines only by functional or metabolic assay, where a transcriptional proxy, such as a named gene set, is required.

**Supplementary Table 1. Definitions used by the four agents and SMART for the source study's key terms.** The paper validates 14 PI3K-AKT pathway genes as hnRNPL targets by RT-qPCR. For glycolysis, proliferation, and migration and invasion, the paper reports functional and metabolic assays only, so no transcriptional definition exists and a proxy is unavoidable.

| | **hnRNPL-regulated PI3K-AKT genes** | **Glycolysis** | **Proliferation; migration and invasion** |
|---|---|---|---|
| **What the paper provides** | 14 RT-qPCR-validated genes (Fig. 4i and Supplementary Fig. 5c of the source study): *COL4A4*, *IRS1*, *GNG12*, *CDK6*, *PIK3CB*, *PCK2*, *F2R*, *PPP2R2C*, *EGFR*, *RXRA*, *ITGA3*, *ITGB1*, *ITGB3*, *ITGB4* | Metabolic assays only | Functional assays only |
| **SMART** | 14/14 validated genes | KEGG_GLYCOLYSIS_ GLUCONEOGENESIS, labeled proxy | Proliferation: HALLMARK_ G2M_CHECKPOINT proposed by SMART when reading the paper; the final definition was the fraction of glioma cells in S or G2/M phase, taken from the cell-cycle labels in the user-supplied metadata of the target dataset. Migration and invasion: HALLMARK_ EPITHELIAL_ MESENCHYMAL_ TRANSITION. All labeled proxy |
| **Biomni** | *PIK3CB*, *EGFR*, *RXRA*, *CDK6*, *IRS1*, *AKT1/2/3*, *MTOR*, *PTEN*, *PIK3CA*, *PIK3R1*; 5/14 overlap | *PFKFB2*, *PKM*, *HK1/2*, *LDHA/B*, *ENO1*, *SLC2A1*, *GAPDH*, *PCK2*; labeled "Glycolysis_paper" and described as curated from the paper text; the paper contains no glycolysis gene set | Not defined; claim not extracted |
| **Claude Code** | *PIK3R1*, *CDK6*, *AKT1/2/3*, *EGFR*, *PIK3CA*, *PIK3CB*, *IRS1*, *PIK3CD*; 4/14 overlap | *GPI*, *HK1*, *ENO1*, *PFKL*, *PKM*, *PFKM*, *LDHA*, *PFKFB2*, *GAPDH*, *SLC2A1*, *HK2*, *SLC2A3*; no reason given | Not defined; claim not extracted |
| **Claude Science** | *CDK6*, *AKT1*, *HIF1A*, *EGFR*, *MTOR*, *AKT2*, *AKT3*, *PIK3CB*, *MYC*, *PIK3CA*, *PTEN*, *ENO1*, *IRS1*, *PFKFB2*, *PKM*, *HK2*, *LDHA*; 4/14 overlap | *HK2*, *PFKFB2*, *PKM*, *LDHA*, *ENO1*, *GAPDH*, *HIF1A*; no reason given | Not defined; claim not extracted |
| **Kosmos** | Hallmark PI3K-AKT-mTOR signaling (105-gene MSigDB set); Kosmos states that the paper supplies no PI3K-AKT signature; 1/14 overlap (*EGFR*) | Hallmark glycolysis (200-gene MSigDB set) | Not defined; claim not extracted |

#### Supplementary Note 2. Comparison of SMART's Results Extraction module with a general-purpose LLM

LLMs summarize biomedical text well[53], but used directly they make small errors when reading a study, such as dropping details, citing figures for the wrong claim, stating results more strongly than the paper supports, or adding content the paper does not contain.

Such errors matter when no one reviews the summary. In an interactive session, a user reads each summary and corrects it before analysis. SMART is an autonomous system and runs without that step, so an uncorrected error passes directly into the analysis that builds on it (Supplementary Note 1).

SMART's Results Extraction module therefore adds a checking step. An LLM agent drafts the summary, and further agents check the draft systematically against the paper's text and figures and correct it before it reaches the user. The agents and their checks are described in Methods.

##### Comparison setup

To illustrate the effect of the checking step, we compared SMART's Results Extraction module against Claude Opus 4.6 used directly through the web interface (Opus 4.6 web). The module is itself built on Claude Opus 4.6, so the two setups share the same base model and any difference between them reflects the design of the module rather than model capability.

The source study is Qin et al.[51], the same paper used in Supplementary Note 1. Opus 4.6 web was given the same article URL and asked for the same three outputs the module produces: clinically relevant findings with figure IDs, mechanistic claims, and definitions of the biological terms used in them.

Both outputs were compared against the source manuscript. Each claim was paired with the paper text supporting it, and each term definition was checked against the relevant text, figure panel or supplementary file. The full evaluation rules, verdict labels and per-item scoring are given in Supplementary Data 6, which also holds the complete output from both setups and the exact prompt used.

##### Comparison results

On this study, SMART's Results Extraction module was correct on all 20 scored items. Opus 4.6 web was correct on 13 of 20, with errors in 2 of 4 clinical findings, 2 of 4 mechanistic claims and 3 of 12 term definitions (Supplementary Data 6, Overview sheet).

#### Supplementary Note 3. Example output of SMART's Data Inspection module

The Data Inspection module, described in Methods, summarizes the structure and metadata of a target dataset. Here it was run on the liver cancer target dataset used throughout this work (GEO accession GSE149614)[54], which contains 71,915 cells from ten hepatocellular carcinoma patients.

Several metadata columns carry the information that downstream modules act on. For example, `celltype` records which of the six major cell populations are present; `site` records where each sample was taken (`Tumor`, `Normal`, `PVTT` or `Lymph`); `stage` and `virus` record cancer stage and viral infection status; and `sample` and `patient` record sample and patient identifiers.

Downstream, the Transferability Assessment module checks the cell types required by the source paper against the `celltype` column. The Task Plan module uses the column names and their values to choose grouping variables and define condition contrasts. Analyses therefore use the dataset's actual column names and categories rather than assumed ones. When a source paper requires finer subtypes than the `celltype` column provides, such as regulatory T cells, the Task Plan module derives them by subclustering (Methods).

The inspection output for the liver cancer dataset is shown below. Its content is unedited; only the font and spacing were adjusted for print.

```
=== COMPREHENSIVE INSPECTION OF LIVER CANCER DATA ===

DATASET OVERVIEW:
• Total cells: 71915
• Total genes: 25712
• Metadata columns: 10

METADATA COLUMNS: orig.ident, nCount_RNA, nFeature_RNA, sample, res.3, site, patient,
stage, virus, celltype

DETAILED COLUMN ANALYSIS:
==============================================================================

--- Column 1: 'orig.ident' ---
• Type: character | Values: 71915 | Missing: 0 (0.0%)
• Unique values: 21
• All unique values: HCC01T, HCC02T, HCC03N, HCC03T, HCC04N, HCC04T, HCC05N, HCC05T,
HCC06N, HCC06T, HCC07N, HCC07P, HCC07T, HCC08N, HCC08P, HCC08T, HCC09N, HCC09T,
HCC10L, HCC10N, HCC10T
• Top categories:
  HCC08T: 4833 (6.7%)
  HCC03T: 4825 (6.7%)
  HCC08N: 4795 (6.7%)
• Sample: HCC01T, HCC01T, HCC01T, HCC01T, HCC01T

--- Column 2: 'nCount_RNA' ---
• Type: numeric | Values: 71915 | Missing: 0 (0.0%)
• Unique values: 23201
• Too many unique values (>60) to list
• Min: 723 | Max: 153832 | Mean: 9811.678 | Median: 4581 | SD: 13065.42
• Q1: 2575 | Q3: 10813
• Sample: 6001, 13864, 3660, 3840, 4102

--- Column 3: 'nFeature_RNA' ---
• Type: integer | Values: 71915 | Missing: 0 (0.0%)
• Unique values: 6702
• Too many unique values (>60) to list
```

• Min: 499 | Max: 7997 | Mean: 2039.377 | Median: 1488 | SD: 1486.838
• Q1: 1018 | Q3: 2525
• Sample: 1780, 2826, 1282, 1227, 1336

--- Column 4: 'sample' ---
• Type: character | Values: 71915 | Missing: 0 (0.0%)
• Unique values: 21
• All unique values: HCC01T, HCC02T, HCC03N, HCC03T, HCC04N, HCC04T, HCC05N, HCC05T, HCC06N, HCC06T, HCC07N, HCC07P, HCC07T, HCC08N, HCC08P, HCC08T, HCC09N, HCC09T, HCC10L, HCC10N, HCC10T
• Top categories:
  HCC08T: 4833 (6.7%)
  HCC03T: 4825 (6.7%)
  HCC08N: 4795 (6.7%)
• Sample: HCC01T, HCC01T, HCC01T, HCC01T, HCC01T

--- Column 5: 'res.3' ---
• Type: integer | Values: 71915 | Missing: 0 (0.0%)
• Unique values: 53
• All unique values: 1, 10, 11, 12, 13, 14, 15, 16, 17, 18, 19, 2, 20, 21, 22, 23, 24, 25, 26, 27, 28, 29, 3, 30, 31, 32, 33, 34, 35, 36, 37, 38, 39, 4, 40, 41, 42, 43, 44, 45, 46, 47, 48, 49, 5, 50, 51, 52, 53, 6, 7, 8, 9
• Min: 1 | Max: 53 | Mean: 17.64 | Median: 15 | SD: 12.752
• Q1: 7 | Q3: 27
• Sample: 13, 16, 25, 2, 2

--- Column 6: 'site' ---
• Type: character | Values: 71915 | Missing: 0 (0.0%)
• Unique values: 4
• All unique values: Lymph, Normal, PVTT, Tumor
• Top categories:
  Tumor: 34414 (47.9%)
  Normal: 28687 (39.9%)
  PVTT: 5971 (8.3%)
• Sample: Tumor, Tumor, Tumor, Tumor, Tumor

--- Column 7: 'patient' ---
• Type: character | Values: 71915 | Missing: 0 (0.0%)
• Unique values: 10
• All unique values: HCC01, HCC02, HCC03, HCC04, HCC05, HCC06, HCC07, HCC08, HCC09, HCC10
• Top categories:
  HCC08: 13770 (19.1%)
  HCC06: 8773 (12.2%)
  HCC10: 8714 (12.1%)
• Sample: HCC01, HCC01, HCC01, HCC01, HCC01

--- Column 8: 'stage' ---
• Type: character | Values: 71915 | Missing: 0 (0.0%)
• Unique values: 5
• All unique values: I, II, IIIA, IIIB, IV
• Top categories:
  IIIB: 19849 (27.6%)
  IIIA: 16782 (23.3%)
  I: 14895 (20.7%)
• Sample: I, I, I, I, I

--- Column 9: 'virus' ---
• Type: character | Values: 71915 | Missing: 0 (0.0%)
• Unique values: 3
• All unique values: HBV, HCV, None
• Top categories:
  HBV: 29734 (41.3%)
  None: 27858 (38.7%)
  HCV: 14323 (19.9%)
• Sample: HBV, HBV, HBV, HBV, HBV

--- Column 10: 'celltype' ---
• Type: character | Values: 71915 | Missing: 0 (0.0%)
• Unique values: 6
• All unique values: B, Endothelial, Fibroblast, Hepatocyte, Myeloid, T/NK
• Top categories:

```
  T/NK: 25591 (35.6%)
  Hepatocyte: 20782 (28.9%)
  Myeloid: 15947 (22.2%)
• Sample: T/NK, Myeloid, T/NK, T/NK, T/NK

==============================================================================
FIRST 5 ROWS:
                        orig.ident nCount_RNA nFeature_RNA sample res.3  site
HCC01T_AAACCTGAGGGCATGT     HCC01T       6001         1780 HCC01T    13 Tumor
HCC01T_AAACCTGAGTCGCCGT     HCC01T      13864         2826 HCC01T    16 Tumor
HCC01T_AAACCTGCATTACCTT     HCC01T       3660         1282 HCC01T    25 Tumor
HCC01T_AAACCTGGTCACACGC     HCC01T       3840         1227 HCC01T     2 Tumor
HCC01T_AAACCTGTCCAGTATG     HCC01T       4102         1336 HCC01T     2 Tumor
                        patient stage virus celltype
HCC01T_AAACCTGAGGGCATGT   HCC01     I   HBV     T/NK
HCC01T_AAACCTGAGTCGCCGT   HCC01     I   HBV  Myeloid
HCC01T_AAACCTGCATTACCTT   HCC01     I   HBV     T/NK
HCC01T_AAACCTGGTCACACGC   HCC01     I   HBV     T/NK
HCC01T_AAACCTGTCCAGTATG   HCC01     I   HBV     T/NK

EXPRESSION DATA:
• Assays: RNA
• Default assay: RNA
• Total UMIs: 705606794 | Mean UMIs/cell: 9811.68 | Mean genes/cell: 2039.38

METADATA STRUCTURE:
'data.frame':71915 obs. of  10 variables:
 $ orig.ident  : chr  "HCC01T" "HCC01T" "HCC01T" "HCC01T" ...
 $ nCount_RNA  : num  6001 13864 3660 3840 4102 ...
 $ nFeature_RNA: int  1780 2826 1282 1227 1336 1553 1616 2059 1518 2385 ...
 $ sample      : chr  "HCC01T" "HCC01T" "HCC01T" "HCC01T" ...
 $ res.3       : int  13 16 25 2 2 38 21 21 2 6 ...
 $ site        : chr  "Tumor" "Tumor" "Tumor" "Tumor" ...
 $ patient     : chr  "HCC01" "HCC01" "HCC01" "HCC01" ...
 $ stage       : chr  "I" "I" "I" "I" ...
 $ virus       : chr  "HBV" "HBV" "HBV" "HBV" ...
 $ celltype    : chr  "T/NK" "Myeloid" "T/NK" "T/NK" ...

SUMMARY STATISTICS:

OBJECT DIMENSIONS:
[1] 25712 71915

=== INSPECTION COMPLETE ===
```

#### Supplementary Note 4. Manual validation of SMART in 10 cases

In this note we describe the manual validation of SMART across 10 cases: one positive control, in which SMART was applied to a source study together with that study's own scRNA-seq dataset, and 9 discovery transfer cases. The complete per-case records are provided in Supplementary Data 7. Supplementary Table 2 records the case identifier, the source paper, the transfer task and the transferability score SMART assigned.

The 9 discovery transfer cases cover the main examples in the main text (ORACLE, UPP1, CYP27A1 and PCSK9), the case used in Supplementary Notes 1 and 2, and one case that SMART judged Infeasible, so that this outcome is also represented. The transferability score is SMART's four-level label; the transfer call is binary, with a study counted as transferring when the score is Middle-high or High.

Each case is recorded in two sheets of Supplementary Data 7. A meta sheet (caseN_meta) holds the source paper DOI, SMART's result extraction for that paper, and the summary of the target scRNA-seq dataset. A validation sheet (caseN_validation) evaluates the SMART workflow end to end, one row per extracted piece: one row per clinically relevant finding, and one row per sentence of the mechanism summary. The Overview sheet of Supplementary Data 7 explains what each column in these tables records.

##### Audit results

Because the paper summary and the data inspection summary are produced once per source study and per target dataset and reused across transfer tasks, as in the large-scale application, these outputs were audited once for each of the 7 source studies and 5 target datasets; all later stages were audited for each case. Of the points in the extracted clinically relevant findings and mechanistic claims, 97% were supported by the source text. The reported cell types and metadata matched the dataset for all 5 target datasets, and the decision to proceed agreed with expert assessment in all 10 cases, including the Infeasible case. In the 9 cases that proceeded, 95% of testable points were addressed by a proper test that SMART executed and reported.

Each biological term the claims depend on was checked against four rules. Where the source paper gives a definition that can be applied in scRNA-seq data, such as a gene set validated by knockdown and RT-qPCR, SMART should use it. Specialized subtypes defined by the paper, such as LRRC15$^{+}$ myofibroblastic cancer-associated fibroblasts, should likewise follow the paper's definition. General cell types, such as T cells or regulatory T cells, should follow the annotation of the target dataset or standard markers. Where the paper defines a term only through functional or metabolic assays, SMART should propose a transcriptional proxy and label it as a proxy. Under these rules, 92% of terms received an appropriate definition. Averaged across the five audit checks (extracted points supported by the source text, term definitions, tests executed and reported, transferability decisions and target dataset summaries), 97% of audited outputs were correct.

**Supplementary Table 2. The 10 manually validated cases.** Case 0 is the positive control, in which the target dataset is the source study's own scRNA-seq dataset; cases 1 to 9 are discovery transfer cases, each pairing one source study with an independent target scRNA-seq dataset. Study IDs refer to Supplementary Table 5; the positive-control study is not among the 121 source studies and is identified by its reference. The last column lists the Supplementary Data file containing each SMART report; for the Infeasible case 9, SMART's output is included in Supplementary Data 7.

| Case | Source study (DOI) | Study ID | Transfer task | Transferability score | Report |
|---|---|---|---|---|---|
| **0** | Qi et al. (colorectal), 10.1038/s41467-022-29366-6 | ref. 22 | colorectal → colorectal (source study's own dataset) | High | Data 8 |
| **1** | Qin et al. (ovarian), 10.1038/s41467-025-60115-7 | S106 | ovarian → glioblastoma | Middle-high | Data 5 |
| **2** | ORACLE (lung), 10.1038/s43018-024-00883-1 | S088 | lung → liver | High | Data 9 |
| **3** | ORACLE (lung), 10.1038/s43018-024-00883-1 | S088 | lung → ovarian | Middle-low | Data 9 |
| **4** | UPP1 (lung), 10.1038/s41467-024-45340-w | S094 | lung → glioblastoma | Middle-high | Data 10 |
| **5** | UPP1 (lung), 10.1038/s41467-024-45340-w | S094 | lung → liver | Middle-high | Data 10 |
| **6** | Das et al. (colorectal), 10.1038/s41586-024-08247-6 | S007 | colorectal → pleural mesothelioma | Middle-high | Data 11 |
| **7** | ORACLE (lung), 10.1038/s43018-024-00883-1 | S088 | lung → pleural mesothelioma | Middle-high | Data 11 |
| **8** | PCSK9 (pancreatic), 10.1038/s41586-025-09017-8 | S115 | pancreatic → ovarian | Middle-high | Data 12 |
| **9** | Heide et al. (ovarian), 10.1038/s41586-025-09303-5 | S101 | ovarian → glioblastoma | Infeasible | Data 7 |

#### Run-to-run stability

Because LLM outputs can vary between runs, each of the 10 cases was run three times independently under the same settings, and the transferability scores were compared across runs. The transfer call was identical across all three runs in all 10 cases, and the transferability score was identical in 7 of 10 (Supplementary Table 3). Scores differed in three cases, each by one level: the positive control, ORACLE transferred to liver cancer and UPP1 transferred to glioblastoma varied between High and Middle-high. No run moved a case across the transfer threshold.

**Supplementary Table 3. Transferability scores across three independent runs of the 10 validated cases.** Cases are numbered as in Supplementary Table 2. 'Same transfer call' indicates whether all three runs agree on transfer (Middle-high or High) versus non-transfer (Middle-low, Low or Infeasible); 'Same transferability score' indicates whether all three runs assign the same score.

| Case | Run 1 | Run 2 | Run 3 | Same transfer call | Same transferability score |
|---|---|---|---|---|---|
| **0** | High | Middle-high | High | Yes | No |
| **1** | Middle-high | Middle-high | Middle-high | Yes | Yes |

| Case | Run 1 | Run 2 | Run 3 | Same transfer call | Same transferability score |
|---|---|---|---|---|---|
| **2** | High | Middle-high | High | Yes | No |
| **3** | Middle-low | Middle-low | Middle-low | Yes | Yes |
| **4** | Middle-high | High | Middle-high | Yes | No |
| **5** | Middle-high | Middle-high | Middle-high | Yes | Yes |
| **6** | Middle-high | Middle-high | Middle-high | Yes | Yes |
| **7** | Middle-high | Middle-high | Middle-high | Yes | Yes |
| **8** | Middle-high | Middle-high | Middle-high | Yes | Yes |
| **9** | Infeasible | Infeasible | Infeasible | Yes | Yes |

#### Decoy test

To test whether SMART simply confirms the claims it is given, we reversed the direction of every finding in the source-study summaries behind the 15 reports featured in this study (Qi et al., ORACLE, UPP1, CYP27A1, PCSK9 and hnRNPL; Methods) and reran these decoys against the same target datasets. All 15 decoy reports were scored Middle-low or Low, including the 12 whose original reports were scored Middle-high or High (Supplementary Table 4).

**Supplementary Table 4. Transferability scores of the original and decoy reports.** For each of the 15 reports featured in this study, the transferability score SMART assigned with the original paper summary and with the decoy summary, in which the direction of every finding was reversed (Methods).

| Report (source study → target) | Original score | Decoy score |
|---|---|---|
| Qi et al. (colorectal) → colorectal | High | Low |
| ORACLE (lung) → colorectal | Middle-high | Middle-low |
| ORACLE (lung) → glioblastoma | Middle-high | Low |
| ORACLE (lung) → liver | High | Middle-low |
| ORACLE (lung) → ovarian | Middle-low | Middle-low |
| ORACLE (lung) → pancreatic | Middle-high | Low |
| ORACLE (lung) → pleural mesothelioma | Middle-high | Low |
| UPP1 (lung) → colorectal | Middle-low | Middle-low |
| UPP1 (lung) → glioblastoma | Middle-high | Low |
| UPP1 (lung) → liver | Middle-high | Low |
| UPP1 (lung) → ovarian | Middle-low | Low |
| UPP1 (lung) → pancreatic | Middle-high | Low |
| CYP27A1 (colorectal) → pleural mesothelioma | Middle-high | Low |
| PCSK9 (pancreatic) → ovarian | Middle-high | Middle-low |
| hnRNPL (ovarian) → glioblastoma | Middle-high | Low |

#### Supplementary Note 5. ORACLE survival association in bulk cohorts

SMART assessed ORACLE as transferable to four of the five non-lung targets (all but ovarian cancer) and to pleural mesothelioma in scRNA-seq data (Supplementary Data 9 and 11). In most of these reports, SMART recommended testing ORACLE against patient survival, which these scRNA-seq datasets do not include. We therefore tested whether the signature is associated with survival in bulk cohorts.

##### Data and scoring

We applied the published ORACLE signature, unmodified, to six TCGA cohorts covering the cancers considered in the transfer analysis: pancreatic (TCGA-PAAD), liver (TCGA-LIHC), glioblastoma (TCGA-GBM), ovarian (TCGA-OV), colorectal (TCGA-COAD and TCGA-READ, pooled) and pleural mesothelioma (TCGA-MESO). Gene expression (RSEM normalized counts, $\log_2(x + 1)$) and overall survival were obtained from the UCSC Xena TCGA hub[45]. For each patient's primary tumor, the ORACLE score was calculated as the sum of the expression of the 23 ORACLE genes weighted by the published coefficients; all 23 genes were present in every cohort. As in the original publications, analyses were restricted to stage I–III; TCGA-GBM, which is uniformly grade IV, was analyzed whole.

##### Statistics

Patients were divided at the within-cohort median into ORACLE high and ORACLE low groups, overall survival was compared with Kaplan–Meier curves and a two-sided log-rank test, and hazard ratios (high versus low) were estimated by univariable Cox regression. To test whether ORACLE's survival association simply reflects proliferation, we compared ORACLE with random 23-gene signatures of the same proliferation content, drawn from proliferation-correlated and other expressed genes and scored with the ORACLE coefficients. Signatures were kept if their score correlated with the Hallmark G2M checkpoint score to within ±0.05 of ORACLE's own correlation, until 1,000 were obtained per cohort. ORACLE and each signature were tested by univariable Cox regression.

##### Results

Patients in the ORACLE high group had significantly shorter overall survival in pancreatic cancer, liver cancer and pleural mesothelioma. No significant association was found in glioblastoma, ovarian or colorectal cancer, with a non-significant trend in the opposite direction in colorectal cancer. Cohort sizes, hazard ratios with 95% confidence intervals and log-rank $P$ values are shown in each panel of Supplementary Fig. 2.

ORACLE was correlated with proliferation (Spearman r with the G2M score 0.65 in liver cancer, 0.51 in pancreatic cancer and 0.70 in mesothelioma). It was more strongly associated with survival than 98% of the matched signatures in liver cancer (empirical $P = 0.021$), 94% in pancreatic cancer ($P = 0.062$) and all of them in mesothelioma ($P = 0.001$).

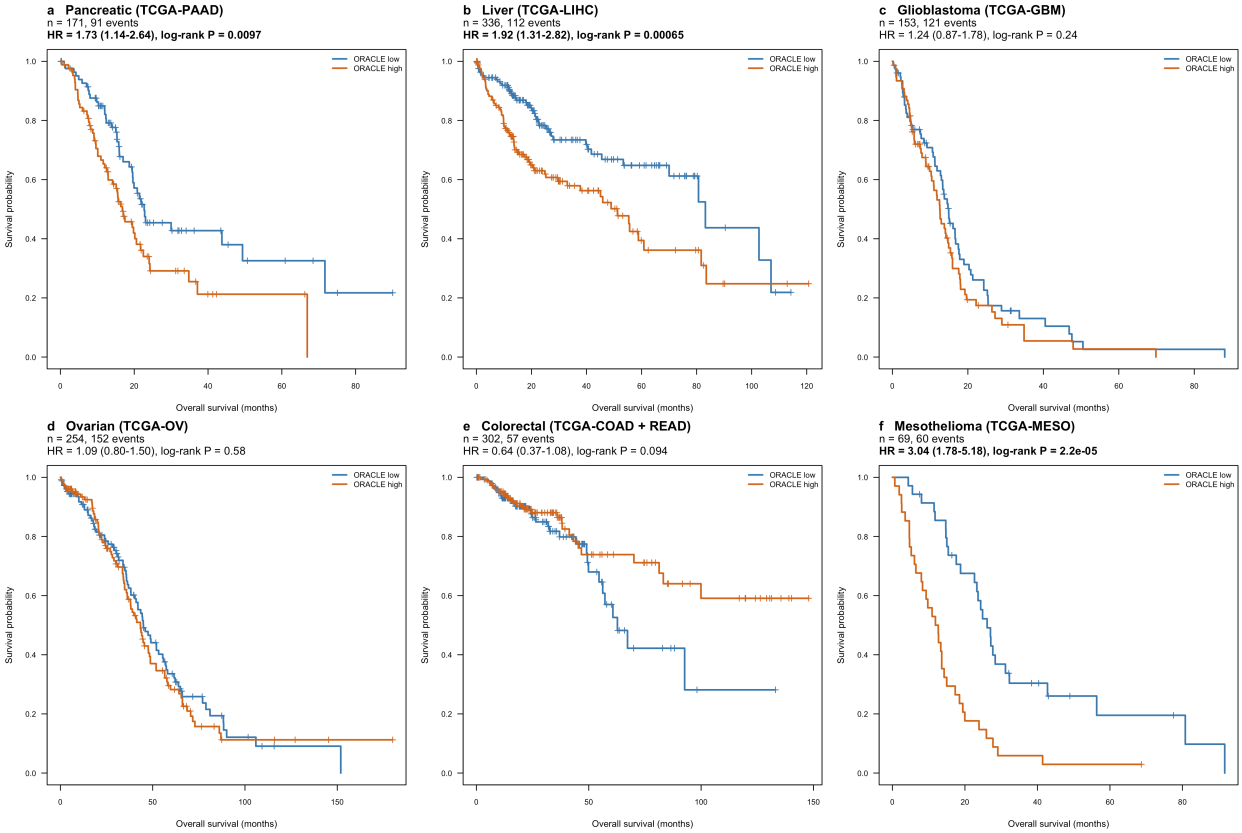


**Supplementary Fig. 2. ORACLE score and overall survival in six TCGA cohorts.** Kaplan–Meier estimates of overall survival for patients dichotomized at the within-cohort median ORACLE score (stage I–III; TCGA-GBM, all patients). Each panel gives the number of patients, the number of deaths, the hazard ratio for ORACLE high versus ORACLE low patients (95% CI, univariable Cox regression) and the *P* value from a two-sided log-rank test.

#### Supplementary Note 6. *PCSK9* expression and GPX4-inhibitor sensitivity in ovarian cancer cell lines

A pancreatic cancer study[43] reported that *PCSK9*-low cells are more sensitive to GPX4 inhibitors than *PCSK9*-high cells. SMART found that the cholesterol biosynthesis arm of this mechanism transferred to ovarian cancer, where *PCSK9*-high tumor cells also showed higher cholesterol biosynthesis activity, and suggested testing GPX4 inhibition (Supplementary Data 12). We therefore tested whether the reported difference in drug sensitivity is seen in ovarian cancer cell lines, using public drug-response and expression data.

##### Data and scoring

Drug-response data were taken from the Cancer Therapeutics Response Portal v2.1 (CTRP)[46,47], downloaded from the NCI CTD² data portal (dataset CTRPv2.1_2016_pub_NatChemBiol_12_109). Sensitivity was measured as the area under the dose–response curve (AUC) published by CTRP, for which a higher value indicates greater resistance. Where a cell line was screened more than once with a compound, the median AUC was used.

Three GPX4 inhibitors were included: ML210, RSL3 (1S,3R-RSL-3) and ML162. The AUC of each compound was z-scored across the ovarian cancer cell lines, and the GPX4-inhibitor resistance score of a line is the mean of its available z-scores. The score is therefore relative among ovarian cancer cell lines.

*PCSK9* expression was taken from DepMap Public 26Q1[48,49] as $\log_2(\text{TPM} + 1)$, using the default expression profile of each cell line. CTRP cell lines were matched to DepMap by cell-line name, and ovarian cancer cell lines were defined by the DepMap lineage annotation (Ovary/Fallopian Tube). Of 39 ovarian cancer cell lines with GPX4-inhibitor data, 38 also had *PCSK9* expression, and these 38 lines were analyzed.

##### Statistics

The 38 lines were divided at their median *PCSK9* expression (0.86 $\log_2(\text{TPM} + 1)$): lines at or below the median were classed as *PCSK9*-low ($n$ = 19) and lines above it as *PCSK9*-high ($n$ = 19). The resistance scores of the two groups were compared with a two-sided Wilcoxon rank-sum test. Spearman's rank correlation between *PCSK9* expression and the resistance score was also computed across all 38 lines. To assess specificity, the same correlation with *PCSK9* expression was computed for the area under the dose–response curve of each of the 407 compounds tested in at least 20 of these lines, and compounds were ranked by correlation.

##### Results

*PCSK9*-low lines were more sensitive to GPX4 inhibitors than *PCSK9*-high lines (median resistance score −0.40 versus 0.11; $P = 0.044$; Fig. 5d), and resistance correlated with *PCSK9* expression across all 38 lines (Spearman $\rho = 0.39$, $P = 0.015$; Supplementary Fig. 3). Among the 407 compounds, ML210, RSL3 and ML162 ranked 19th, 27th and 42nd (top 4.7%, 6.6% and 10.3%).

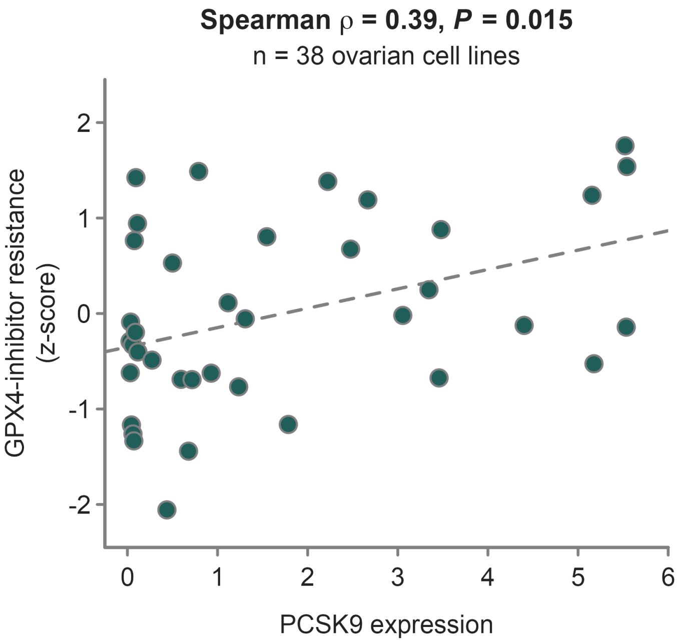


**Supplementary Fig. 3. Correlation between *PCSK9* expression and GPX4-inhibitor resistance across 38 ovarian cancer cell lines.** Each point is one of the 38 ovarian cancer cell lines analyzed in this Note. *PCSK9* expression is $\log_2$(TPM + 1) from DepMap; GPX4-inhibitor resistance is the mean of the z-scored CTRP areas under the dose–response curve for ML210, RSL3 and ML162. The dashed line is a least-squares linear fit, shown as a guide. ρ and P, Spearman's rank correlation.

#### Supplementary Table legends

*Legends for Supplementary Tables 1 to 4 are given with the tables, in Supplementary Notes 1 and 4.*

**Supplementary Table 5. Source studies and transferability scores for all 726 SMART reports.** The 121 single-cell RNA-sequencing studies used as sources (17 colorectal, 24 glioblastoma, 29 liver, 24 lung, 12 ovarian and 15 pancreatic cancer), published in 2024–2025 in Nature, Nature Cancer, Nature Communications, Nature Genetics, Nature Immunology or Nature Medicine, with title, journal, year, DOI and the transferability score SMART assigned in each of the six target datasets (Methods).

**Supplementary Table 6. Mechanistic claims involving *MYC* and their transfer to glioblastoma.** The 17 mechanistic claims involving *MYC* extracted from non-glioblastoma source studies, with the source study for each, the direction of the claim, whether it transferred to glioblastoma, and, for claims already documented in glioblastoma, the supporting publication.

**Supplementary Table 7. Topic assignment of the ovarian and pancreatic source studies.** The 27 ovarian and pancreatic source studies grouped in Fig. 5c, each assigned by manual reading to one topic (lipid metabolism, fibroblasts, immune environment or tumor cell-intrinsic programs). Study identifiers and transferability scores are given in Supplementary Table 5.

#### Supplementary Data legends

Differential expression is computed with the two group labels in alphabetical order, the first serving as the reference; for a comparison of A-high and A-low cells, log fold changes therefore give A-low relative to A-high, and a negative value indicates higher expression in A-high cells. SMART's volcano plots label the direction on the x axis, and its enrichment plots state in the title the group in which the genes are upregulated.

**Supplementary Data 1 to 4. Outputs of the comparator agents.** Complete outputs of Biomni (Supplementary Data 1), Claude Code (Supplementary Data 2), Claude Science (Supplementary Data 3) and Kosmos (Supplementary Data 4) on the evaluation task described in Supplementary Note 1. Biomni and Kosmos returned PDF reports directly; Claude Code and Claude Science returned Markdown, which we converted to PDF without altering content.

**Supplementary Data 5. SMART report for the example in Supplementary Note 1.** The complete SMART report for the ovarian cancer study of Qin et al. evaluated against the glioblastoma single-cell dataset, which is SMART's output in the comparisons of Supplementary Notes 1 and 2 and case 1 of Supplementary Note 4.

**Supplementary Data 6. Comparison of SMART's Results Extraction module with a general-purpose LLM.** Clinically relevant findings, mechanistic claims and term definitions extracted by SMART's Results Extraction module for the study in Supplementary Note 2, beside those from Claude Opus 4.6 used through its web interface (Opus 4.6 web) given the same article URL. The Overview sheet gives a summary of scores, the evaluation rules and the definition of each error label; the Meta sheet gives both outputs in full and the prompt; the Clinical, Mechanism and Definition sheets give per-item scoring.

**Supplementary Data 7. Manual audit of the SMART workflow in the 10 validated cases.** For the positive control and each of the 9 discovery transfer cases of Supplementary Note 4, the audit of SMART's outputs against the source manuscript and the target dataset, covering the extracted findings and claims, the term definitions and the tests reported for each claim. A meta sheet (caseN_meta) holds the source paper DOI, SMART's result extraction and the summary of the target dataset; a validation sheet (caseN_validation) evaluates the workflow end to end, one row per extracted piece. The Overview sheet explains each column. For the Infeasible case 9, the sheets also hold SMART's Transferability Assessment output.

**Supplementary Data 8. SMART report for the positive control.** The complete SMART report for the colorectal cancer study of Qi et al. paired with that study's own single-cell dataset, discussed in Fig. 2 and recorded as case 0 in Supplementary Note 4.

**Supplementary Data 9. SMART reports on the transfer of ORACLE.** Complete reports for the transfer of the ORACLE prognostic signature from lung cancer to colorectal, liver, ovarian and pancreatic cancer and glioblastoma; the reports for liver and ovarian cancer are cases 2 and 3 of Supplementary Note 4.

**Supplementary Data 10. SMART reports on the transfer of UPP1.** Complete reports for the transfer of the UPP1 study from lung cancer to colorectal, liver, ovarian and pancreatic cancer and glioblastoma; the reports for glioblastoma and liver cancer are cases 4 and 5 of Supplementary Note 4.

**Supplementary Data 11. SMART reports on the transfer of CYP27A1 and ORACLE to pleural mesothelioma.** Complete reports for the transfer of the CYP27A1 study from colorectal cancer, and of the ORACLE signature from lung cancer, to a pleural mesothelioma single-cell dataset; these are cases 6 and 7 of Supplementary Note 4.

**Supplementary Data 12. SMART report on the transfer of PCSK9.** The complete SMART report for the transfer of the PCSK9 study from pancreatic to ovarian cancer; this is case 8 of Supplementary Note 4.
